# Antagonistic kinesin-14s within a single chromosomal drive haplotype

**DOI:** 10.1101/2025.02.05.636711

**Authors:** Meghan J. Brady, Anjali Gupta, Jonathan I. Gent, Kyle W. Swentowsky, Robert L. Unckless, R. Kelly Dawe

## Abstract

In maize, there are two meiotic drive systems that operate on large tandem repeat arrays called knobs that are found on chromosome arms. One meiotic drive haplotype, Abnormal chromosome 10 (Ab10), encodes two kinesin proteins that interact with two distinct tandem repeat arrays in a sequence-specific manner to confer meiotic drive. The kinesin KINDR associates with knob180 repeats while the kinesin TRKIN associates with TR-1 repeats. Prior data show that meiotic drive is conferred primarily by the KINDR/knob180 system, with the TRKIN/TR-1 system having little or no role. The second meiotic drive haplotype, K10L2, shows low levels of meiotic drive and only encodes the TRKIN/TR-1 system. Here we used long-read sequencing to assemble the K10L2 haplotype and showed that it has strong homology to an internal portion of the Ab10 haplotype. We also carried out CRISPR mutagenesis of *Trkin* to test the role of *Trkin* on Ab10 and K10L2. The data indicate that the *Trkin* gene on Ab10 does not improve drive or fitness but instead has a weak deleterious effect when paired with a normal chromosome 10. The deleterious effect is more severe when Ab10 is paired with K10L2: in this context functional *Trkin* on either chromosome nearly abolishes Ab10 drive. We modeled the effect of *Trkin* on Ab10 and found it should not persist in the population. We conclude that *Trkin* either confers an advantage to Ab10 in untested circumstances or that it is in the process of being purged from the Ab10 population.

**ARTICLE SUMMARY:** Mendel’s first law states that paired chromosomes are transmitted through meiosis at equal frequencies. Some chromosome variants, however, are transmitted at higher frequencies in a process called meiotic drive. We wanted to know the function of a motor protein called TRKIN that is encoded on a maize meiotic drive chromosome. Surprisingly, we found that TRKIN provides no advantage to the meiotic driver and instead seems to be deleterious, suggesting it had a function in a wild ancestor but is now being purged from the population. The results illustrate how genomes are shaped by often-conflicting forces of selection and selfish genetic elements.

## INTRODUCTION

Selfish genetic elements (i.e. transmission ratio distorters) are structural elements of the genome that increase their own representation in the next generation despite conferring no fitness advantage (Burt and Trivers 2008). Meiotic drivers, one class of selfish genetic element, gain their advantage by altering meiosis so that they are transmitted to more than 50% of the gametes (Lindholm *et al*. 2016). Examples of meiotic drive that operate at the level of meiosis are centromere drive, where larger centromeres are preferentially transmitted over smaller centromeres (Fishman and Kelly 2015; Lampson and Black 2017; Clark and Akera 2021; Dawe 2022), the segregation of certain B chromosomes (Fishman and Kelly 2015; Lampson and Black 2017; Clark and Akera 2021; Dawe 2022), and the maize Abnormal chromosome 10 haplotype (Ab10) (Fishman and Kelly 2015; Lampson and Black 2017; Clark and Akera 2021; Dawe 2022). There are also many other examples of drivers that exhibit preferential transmission but gain their advantage outside of meiosis (Lindholm *et al*. 2016). Selfish genetic elements are implicated in critical evolutionary processes such as extinction, speciation, recombination, and genome size evolution (Agren and Clark 2018). Ab10 is of particular interest as it has had a significant impact on shaping the evolution of maize, one of the most economically important crops (Buckler *et al*. 1999).

As much as >15% of the maize genome is composed of tandem repeat arrays (Hufford *et al*. 2021). One form of tandem repeat is referred to as knobs, and come in two different sequence classes, TR-1 and knob180. The Ab10 meiotic drive haplotype contains long arrays of both knob repeats as well as two kinesin protein-encoding genes: *Kindr* and *Trkin*. KINDR physically associates with knob180 knobs and TRKIN associates with TR-1 knobs (Figure 1a). Both kinesins pull their respective knobs ahead of the centromere during meiotic anaphase to cause their preferential transmission to the egg cell during female meiosis (Dawe 2022) (Figure 1b). Knobs throughout the genome are also preferentially transmitted when Ab10 is present. Both knob180 and TR-1 are conserved and abundant across the *Zea* genus and in *Tripsacum dactyloides* suggesting that Ab10 may have originated deep in the evolutionary history of the grass family (Buckler *et al*. 1999; Swentowsky *et al*. 2020). The KINDR/knob180 system is primarily responsible for the preferential transmission of Ab10 while the TRKIN/TR-1 system contributes little, if at all (Kanizay *et al*. 2013a; Dawe *et al*. 2018). Nevertheless, *Trkin* is present on multiple Ab10 haplotypes in both teosinte and maize suggesting it may have been maintained via selection over the ∼8700 years since their divergence (Piperno *et al*. 2009; Higgins *et al*. 2018; Swentowsky *et al*. 2020).

**Figure 1:**
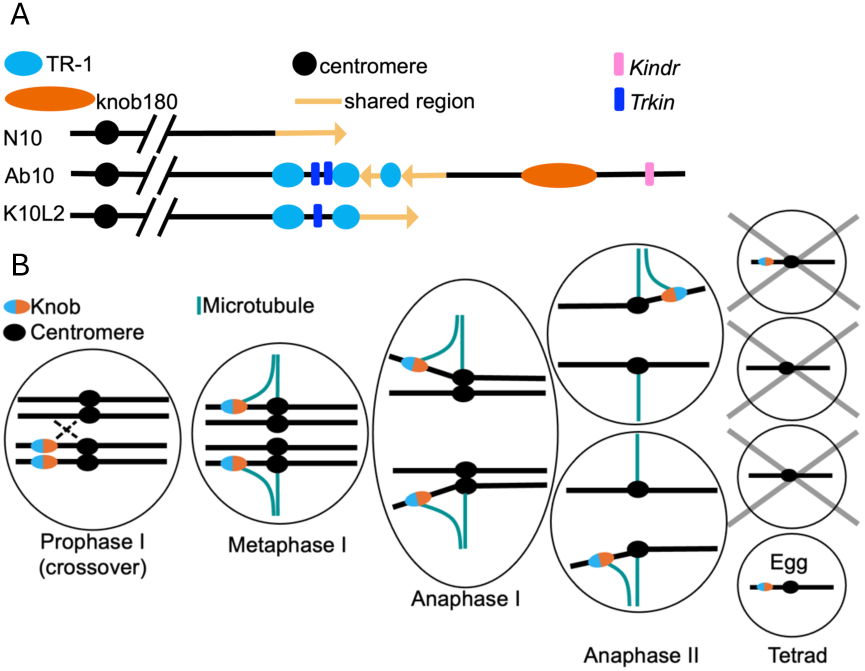
Diagram of Maize Chromosome 10 Haplotypes. A. Diagram of the structure of three chromosome 10 haplotypes. The orientation of the shared region on K10L2 was unknown prior to this study, the orientation we determined is shown. B. Model of Ab10 meiotic drive. For Ab10 drive to occur during female meiosis, the plant must be heterozygous for Ab10. Then recombination must occur between the centromere and the beginning of the Ab10 haplotype. During metaphase TRKIN associates with TR-1 knobs and KINDR associates with knob180 knobs. Both kinesin-14 proteins then drag the knobs ahead of the centromere during anaphase I and II causing their preferential transmission to the top and bottom cells of the meiotic tetrad. Since only the bottom-most cell becomes the egg cell, Ab10 is overrepresented in progeny (Dawe *et al*. 2018; Swentowsky *et al*. 2020).

K10L2 is a structurally and functionally distinct variant of chromosome 10 that expresses TRKIN during meiosis and activates neocentromeres at TR-1 repeats (Kanizay *et al*. 2013a) (Figure 1). K10L2 demonstrates weak (1-2%) but statistically-significant meiotic drive (Kanizay *et al*. 2013a). Additionally, it has been identified in at least 12 disparate maize landrace populations suggesting it may be an important part of the Ab10 system (Kanizay *et al*. 2013a). One to two percent drive should be sufficient to cause K10L2 to rapidly spread throughout a population as long as it isn’t associated with negative fitness consequences (Hartl 1970). K10L2 is also a very effective competitor against Ab10. When Ab10 is paired with K10L2, Ab10 drive is almost completely suppressed (Kanizay *et al*. 2013a). It has been speculated that both the drive of K10L2 and the suppressive effect of K10L2 on Ab10 are mediated by the TRKIN/TR-1 system (Swentowsky *et al*. 2020).

The fitness costs commonly imposed on the genome by selfish genetic elements selects for suppressors throughout the genome (Price *et al*. 2020). In the Ab10 system, K10L2 and N10 both represent disadvantaged loci. K10L2 can be thought of as both a disadvantaged locus carrying a highly effective suppressor when interacting with Ab10 and as an independent driver when interacting with N10. Both Ab10 and K10L2 have what appear to be suppressors of the KINDR/knob180 drive system. N10 carries a pseudo-*Kindr* locus that produces siRNAs that may suppress *Kindr* expression and reduce drive (Dawe *et al*. 2018). K10L2 also acts as a suppressor of Ab10 with the likely mechanism being the TRKIN/TR-1 drive system. The evolution of suppressors by co-opting the machinery of drive has been observed before (Price *et al*. 2020). For example, the *wtf* genes in *Schizosaccharomyces pombe* represent a toxin-antidote system. There are *wtf* loci carrying only the antidote that behave as suppressors to intact *wtf* loci (Bravo Núñez, María Angélica, Lange, Jeffrey J, Zanders, Sarah E 2018). If the Ab10 drive system followed the same model, we would expect that the TRKIN/TR-1 system (i.e. a suppressor) would appear only on K10L2 or N10. How or why *Trkin* persists on Ab10 while conferring no apparent benefit in terms of drive, and likely contributing to the suppression of drive when paired with K10L2, is unclear.

Several hypotheses have been proposed to resolve the conundrum of the *TRKIN/TR-1* drive system on Ab10, all suggesting that *Trkin* improves the fitness of Ab10. The main ideas are that *Trkin* may: 1) increase Ab10 drive or 2) reduce the negative fitness effects associated with Ab10 (Swentowsky *et al*. 2020). In previous work the favored hypothesis was that *Trkin* reduces meiotic errors caused by the rapid movement of knobs during meiotic anaphase (Swentowsky *et al*. 2020). In this study, we set out to determine what effect *Trkin* has on Ab10 that may help to explain its persistence. We assembled the K10L2 haplotype and compared it to Ab10, then conducted drive and fitness assays of K10L2 and Ab10 haplotypes carrying *trkin* null alleles. Finally, we used mathematical modeling to better understand the predicted population dynamics of Ab10 haplotypes that carry *Trkin*.

## RESULTS

### Assembly of K10L2 and Ab10

We began by generating a new assembly of Ab10 using PacBio HiFi. The Ab10 haplotype has been challenging to accurately assemble due to the prevalence of multiple repetitive arrays (i.e. knobs) that are notoriously difficult to assemble (Tørresen *et al*. 2019). The previous assembly of B73-Ab10 v1 was conducted with PacBio CLR data (single long reads) which have a higher error rate (Liu *et al*. 2020; Hon *et al*. 2020). To assess the quality and fidelity of the new assembly, we compared sequence homology between B73-Ab10 v1 (Liu *et al*. 2020) and the new assembly, B73-Ab10 v2. We found strong homology between the assemblies and the same relationship to N10 as previously reported (Supplementary Figure 1, Supplementary Figure 2b). In both assemblies the Ab10 haplotype is located at the end of the long arm of chromosome 10 as expected (Liu *et al*. 2020; Dawe 2022). The total size is unknown because of N-gaps predominantly within tandem-repeat arrays but, using the B73-Ab10 v2 assembly, we estimate the Ab10 haplotype contains about 77 Mb of sequence, with the proximal edge traditionally defined as the *colored1* (*r1)* gene (a linked marker used to track Ab10 in crosses). We identified two large inverted segments homologous to N10 within the haplotype of 4.8 Mb and 9.5 Mb respectively (shared region) (Figure 1a, Supplementary Figure 2b). These are slightly longer than reported in B73-Ab10 v1 assembly (Liu *et al*. 2020). There are three TR-1 knobs (assembled length=8.7 Mb collectively) and a very large knob180 knob (partially assembled length=8.5 Mb). Both the TR-1 and knob180 knobs assembled lengths are slightly lower than in the B73-Ab10 v1 assembly (Liu *et al*. 2020). Using data from terminal deletion lines of Ab10 in a different inbred background, we determined that the Ab10 knob is ∼30.67 Mb long indicating it is only 28% assembled (Brady *et al*. 2024). There is also at least ∼22 Mb of sequence that is unique to Ab10. The 1.8 Mb region between the first two TR-1 knobs includes two copies of *Trkin* (Figure 2). The region to the right of the large knob180 knob contains an array of *Kindr* genes. Interestingly, there was a marked reduction in percent identity between the two assemblies over large tandem arrays like *Kindr* (Supplementary Figure 1). This is likely due to the increased accuracy of PacBio HiFi reads (Hon *et al*. 2020). In fact, we identified 10 copies of *Kindr* in B73-Ab10 v2 instead of 9 as previously reported in B73-Ab10 v1 (Liu *et al*. 2020) (Supplementary Figure 2d).

**Figure 2:**
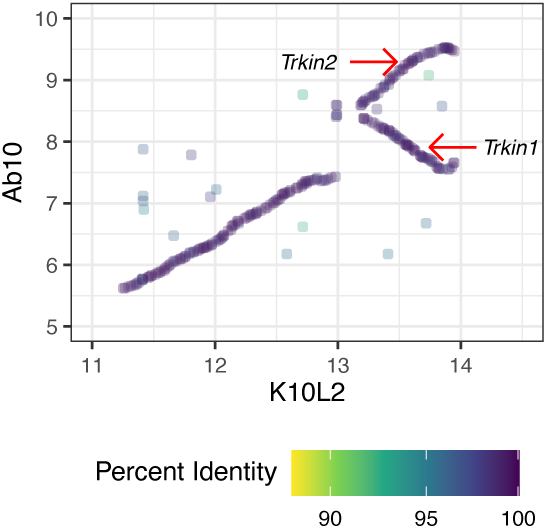
Sequence Comparison of *Trkin* Bearing Region on Ab10 and K10L2. Each dot marks the start of a maximal unique match (MUM) of at least 300bp long between the Ab10 and K10L2 haplotype, which begin at the *colored1* gene (Marçais *et al*. 2018). Coordinates start at the *colored1* gene. The color of each dot represents the percent identity of that match. All large knob arrays were removed for the sake of clarity. Both Ab10 *Trkin* genes are marked. The K10L2 and Ab10 assemblies refer to the assemblies generated in this work.

We next assembled the K10L2 haplotype. We found a distinct structure with two large TR-1 knobs (15.5 Mb collectively) and a 2.7 Mb non-shared region with a single copy of *Trkin* between them (Figure 1a, non-shared means a lack of homology to N10). Otherwise, we found no large inversions or other rearrangements relative to N10 (Supplementary Figure 2a). Additionally, we found no tandemly repeated genes (i.e. *Kindr* array), which are common on Ab10 (Supplementary Figure 2c,d) (Dawe *et al*. 2018). Sequence comparisons revealed the region between the two TR-1 knobs on K10L2 has strong homology to the *Trkin* bearing region on Ab10. However, unlike K10L2, Ab10 contains an inverted duplication with a second copy of *Trkin* (Figure 2, Supplementary Figure 2e) (Swentowsky *et al*. 2020). The second copy of *Trkin* on Ab10 was previously thought to be a pseudogene and was referred to as Ab10 *pseudo*-*Trkin1* (Swentowsky *et al*. 2020). During this study we found that the coding sequence of *pseudo*-*Trkin1* was misinterpreted, and that it instead encodes a full-length open reading frame. Accordingly, we have renamed *pseudo*-*Trkin1* to *Trkin2* (Figure 3).

**Figure 3:**
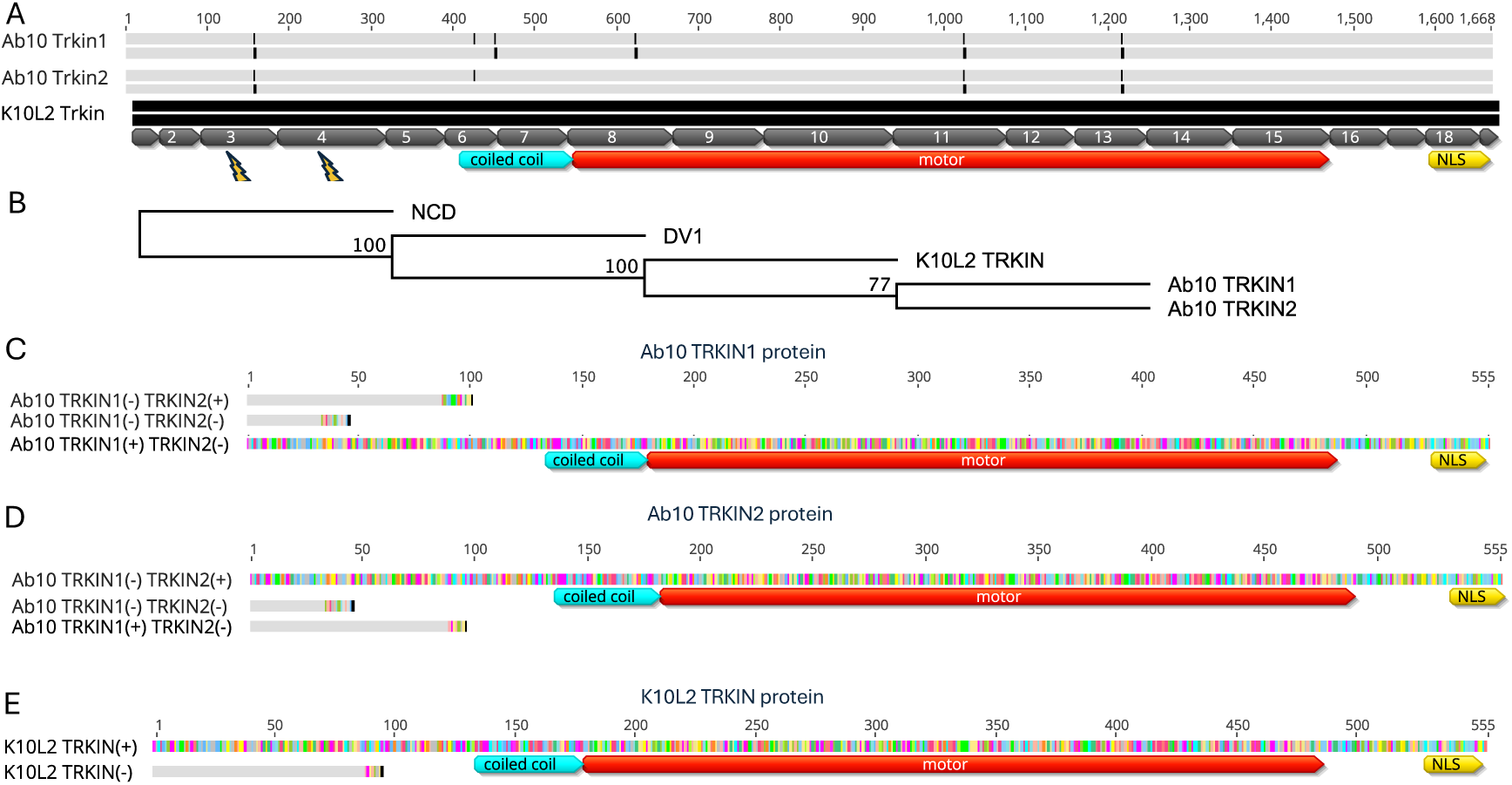
Comparison of *trkin* and Mutants. A. A coding sequence alignment (top bar) and protein translation (bottom bar) of all three *Trkin* sequences. Grey indicates sequence that is identical to the K10L2 *Trkin*, black indicates sequence that is different from the K10L2 *Trkin*. Exon boundaries are marked by numbered grey boxes. Protein domains are marked by colored boxes and labeled by domain type. NLS = nuclear localization signal (Swentowsky *et al*. 2020). Lightning bolts indicate exons that Cas9 was targeted to. B. Neighbor joining consensus tree using Jukes-Cantor model and 1000 bootstraps of protein motor domain for all TRKIN alleles, the most closely related *Zea mays* gene *Dv1*, and the *Drosophila melanogaster* Ncd gene as an outgroup (Swentowsky *et al*. 2020). Number at nodes indicate the number of replicate trees supporting that node. C. Ab10 *Trkin*1 protein alignment. Grey indicates sequence identical to the intact (+) Ab10 *Trkin1*. Color indicates sequence that is different from the intact (+) Ab10 *Trkin1*. Ab10 *trkin1(-) Trkin2(+)* and Ab10 *trkin1(-) trkin2(-)* are truncated as a result of stop codons. D. Ab10 TRKIN2 protein alignment. Grey indicates sequence identical to the intact (+) Ab10 TRKIN2. Color indicates sequence that is different from the intact (+) Ab10 TRKIN2. Ab10 TRKIN1*(+)* TRKIN2*(-)* and Ab10 TRKIN1*(-)* TRKIN2*(-)* are truncated as a result of the introduction of a stop codon. E. K10L2 TRKIN protein alignment. Grey indicates sequence identical to the intact (+) K10L2 <u>TRKIN</u>. Color indicates sequence that is different from the intact (+) K10L2 TRKIN. K10L2 TRKIN(-) is truncated as a result of the introduction of a stop codon. C, D, E. Protein domains are marked by colored boxes labeled by domain type. NLS = nuclear localization signal (Swentowsky *et al*. 2020).

### Genomic sequence of three *Trkin* genes reveals near identical intronic transposons

We annotated the K10L2 and new B73-Ab10 v2 assemblies using BRAKER v3.0.8 (Gabriel *et al*. 2024), which was not available at the time of the B73-Ab10 v1 assembly (Liu *et al*. 2020). This allowed us to identify the full unbiased structure of each independent copy of *Trkin* on both Ab10 and K10L2. In line with the strong homology between the K10L2 haplotype and Ab10, inspection of the *Trkin* genomic sequence revealed a similar atypical structure between all three *Trkin* genes. Ab10 *Trkin1* spans 113 Kb and Ab10 *Trkin2* spans 99 Kb, while the K10L2 *Trkin* spans 89 Kb. The size differences are due to the presence of nine transposable elements in the introns of Ab10 *Trkin1* and *two* transposable elements in the introns of Ab10 *Trkin2* relative to K10L2 *Trkin*. The transposable elements in Ab10 *Trkin1* and Ab10 *Trkin2* are not shared suggesting duplication and divergence after separation from the K10L2 *Trkin*. Notably, Ab10 *Trkin1* and *Trkin2* carry all the transposable elements that are present in K10L2 *Trkin* (Figure 4). These data suggest that K10L2 *Trkin* is ancestral to the Ab10 *Trkin* genes.

**Figure 4:**
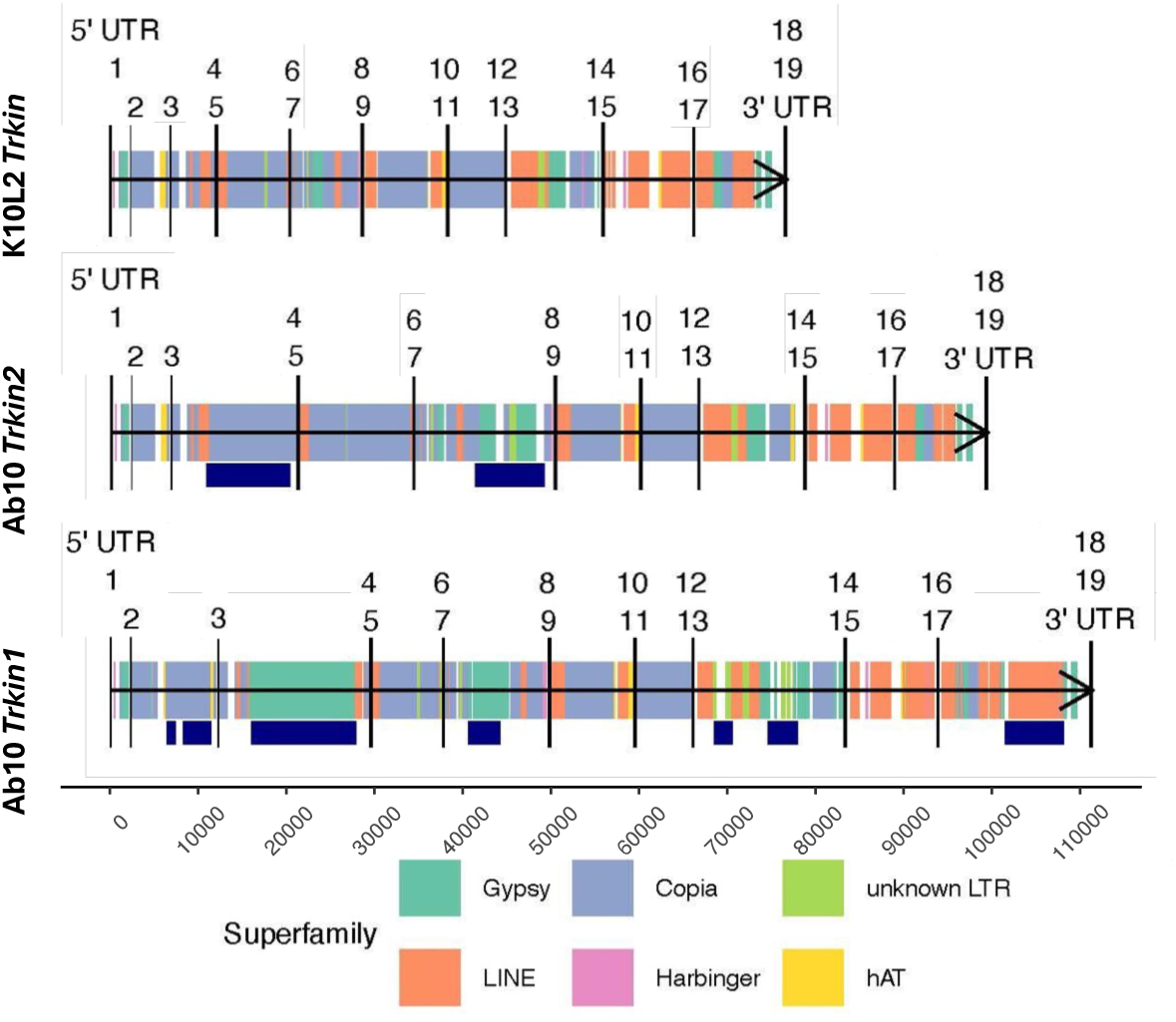
Comparison of Transposable Element (TE) Composition Between All *Trkin* genes. Genomic sequences for all three *Trkin* alleles, represented by a horizontal black line, are shown from Ab10 and K10L2. Vertical long black lines indicate *Trkin* exons. Short colored boxes centered on the horizontal black line indicate annotated transposable elements colored by their superfamily. Navy bars below the annotated TE blocks indicate insertions unique to that *Trkin* allele.

### Comparison of three *Trkin* CDS sequences reveals very few differences

Interrogation of the *Trkin* annotated coding sequence revealed that all three *Trkin* genes are remarkably similar with no significant evidence of functional divergence (Figure 3a). The K10L2 *Trkin* CDS contains six point mutations relative to Ab10 *Trkin1*. Five of these produce nonsynonymous amino acid substitutions (one in an unstructured region, one in the coiled coil domain, and three in the motor domain). The K10L2 *Trkin* CDS contains only four point mutations relative to Ab10 *Trkin2*, of which three cause nonsynonymous amino acid substitutions (one in an unstructured region and two in the motor domain). Ab10 *Trkin1* and Ab10 *Trkin2* differ by only two point mutations resulting in non-synonymous amino acid substitutions (one in the coiled coil domain and one in the motor domain) (Figure 3a). These data suggest that differing effects of *Trkin* between Ab10 and K10L2, if any exist, are not due to differences in the protein itself.

We next wondered what the relationship between the three *Trkin* genes might be. We generated a neighbor joining tree using the amino acids of the motor domain of all three *Trkin* gene as well as their most similar maize gene as an outgroup. We found that Ab10 *Trkin1* and Ab10 *Trkin2* are more similar to each other than to K10L2 *Trkin* (Figure 3b). This relationship suggests that the Ab10 *Trkin* genes duplicated after they diverged from K10L2 *Trkin*, in agreement with the inferences from the TE profile (Figure 4).

### Gene orthology between three chromosome 10 haplotypes finds high agreement in the *Trkin* bearing region and unexpected orthologs in the Ab10 non-shared region

We next investigated the gene orthology between all three assembled structural variants of chromosome 10 (Figure 5). We define the shared regions of both K10L2 and Ab10 as the regions with significant homology to N10, and the non-shared regions as the regions without significant homology to N10 (Supplementary Figure 2, Figure 5). We found that there were 12 gene ortholog pairs between the Ab10 *Trkin* region and K10L2 *Trkin* region representing 44% (12/27) of annotated genes in this region on K10L2 and 66% (12/18) of the annotated genes in this region of Ab10 (Supplementary Table 1, Supplementary Table 2, Supplementary Table 3, Figure 5). There were also unexpected gene ortholog pairs particularly between the shared region of K10L2 and the non-shared region of Ab10 (Supplementary Table 4, Figure 5). Interestingly, using our new annotations, we identified 10 previously unknown gene orthologs between N10 and Ab10 in the non-shared region (Supplementary Table 5, Figure 5). Among the newly identified genes are nine partial copies of a gene homologous to *nrpd2*/*e2*, which is related to RNA dependent DNA methylation (Figure 5, Supplementary Figure 3). This is of particular interest as it has been hypothesized that RNA dependent DNA methylation may be related to the antagonistic dynamics between Ab10 and the host genome (Dawe *et al*. 2018).

**Figure 5:**
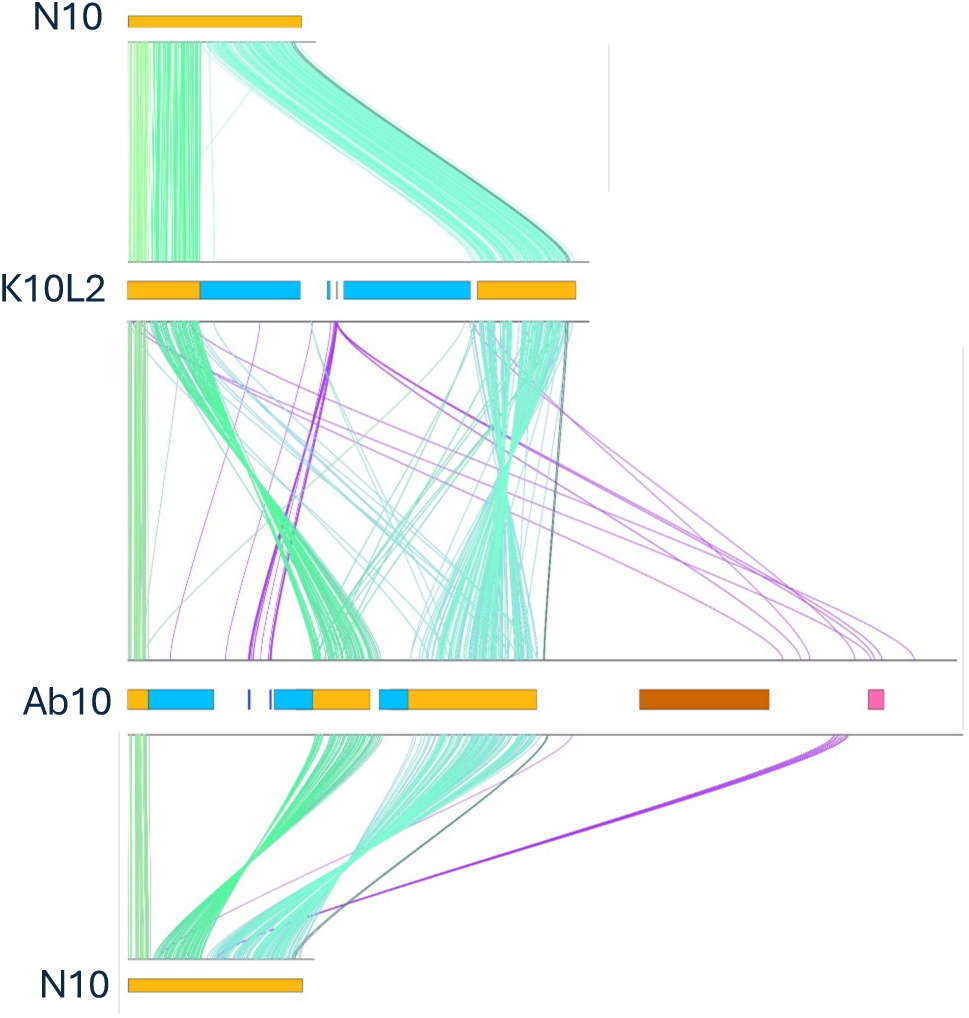
Gene Ortholog Comparisons Among Chromosome 10 Haplotypes. Each line represents a gene ortholog pair as determined by OrthoFinder (Emms and Kelly 2019). Shades of green represent gene ortholog pairs in the shared region. Purple represents gene ortholog pairs outside of the shared region. Relevant regions of each haplotype are marked by colored bars: gold = shared, light blue = TR1 knob, dark blue = *Trkin*, dark orange = knob180 knob, pink = *Kindr*. K10L2 and Ab10 refer to the assemblies generated in this work. N10 refers to the B73 v5 assembly (Hufford *et al*. 2021).

### Ab10 non-shared region annotations are enriched for RNA dependent DNA methylation GO terms

We went on to perform a functional annotation of the Ab10 and K10L2 haplotypes using EnTAP (Supplementary Table 1, Supplementary Table 2) (Hart *et al*. 2020; Gabriel *et al*. 2024). Incorporating all gene annotations, Ab10 is significantly enriched for GO terms related to RNA dependent DNA methylation (Supplementary Figure 4), a result that that likely reflects the high copy number of *nrpd2*/*e2*. We also reduced all known tandemly duplicated genes to a single copy and reran the analysis. Under these circumstances, Ab10 is enriched for GO terms related to meiotic organization and microtubule based movement in agreement with our understanding of the mechanism (Supplementary Figure 5) (Dawe 2022). Ab10 is enriched for RNA dependent DNA methylation when considering gene copy number, but not when considering only unique genes. In contrast, the K10L2 region was only significantly enriched for general reproductive processes, ATP hydrolysis, and several other miscellaneous GO terms (Supplementary Figure 6).

### *Trkin* expression in K10L2 and Ab10 lines

The *Trkin* copy number difference between Ab10 and K10L2 led us to wonder if they may also have expression level differences. We obtained RNA sequencing for Ab10 and K10L2 and mapped it to the B73-Ab10 v1 assembly (Liu *et al*. 2020; Swentowsky *et al*. 2020). The data revealed no consistent difference in *Trkin* expression between Ab10 bearing two copies and K10L2 bearing one copy of *Trkin* (Supplementary Figure 7).

We also assessed the relative expression levels of *Trkin1* and *Trkin2* on Ab10. Analysis of RNA-seq data from ten tissues from a homozygous Ab10 line (Liu *et al*. 2020) indicated that the expression of *Trkin2* is ∼93% lower on average than *Trkin1* (t = 6.5, df = 41.4, p-value = 6e-08) (Supplementary Figure 8).

### Generation of *trkin* knockout mutants on K10L2 and Ab10

To knock out the *trkin* gene on both K10L2 and Ab10, we designed a CRISPR construct with three guide RNAs targeting exon 3 and exon 4 of the *Trkin* gene (Figure 3). When we initiated the CRISPR mutagenesis, we were under the impression that Ab10 *Trkin2* was a pseudogene, and did not assay it for mutations; the primers were designed to be specific to Ab10 *Trkin1* (Supplementary table 6) (Swentowsky *et al*. 2020). Later, when we determined that Ab10 *Trkin2* is likely functional, we developed primers specific to Ab10 *Trkin2* and found that it is mutated in the line we were using as a positive control (Supplementary Table 6, Figure 3d). We isolated the following mutations: K10L2 *Trkin(+),* K10L2 *trkin(-)*, Ab10 *Trkin1(+) trkin2(-)*, Ab10 *trkin1(-) Trkin2(+)*, and Ab10 *trkin1(-) trkin2(-)* (Figure 3c,d,e). For K10L2, we had both a true wild type and a *trkin* mutant. For Ab10, we lacked a true wild type, so compared lines carrying either *Trkin1* or *Trkin2* alone to double mutants lacking both *trkin1* and *trkin2*.

Based on the strong correlation between *Trkin* and TR-1 neocentromere activity (Swentowsky *et al*. 2020), we expected *trkin* mutants to lack TRKIN protein and visible TR-1 neocentromeres at meiosis. In the Ab10 *trkin1*(-) *trkin2*(-) double mutant plants we could not detect TRKIN by immunostaining and observed no TR-1 neocentromeres by FISH (Figure 6, Figure 7), whereas Ab10 *Trkin1*(+) *Trkin2(-)* showed strong TRKIN staining and TR-1 neocentromeres (Figure 6, Figure 7). In the K10L2 *trkin*(-) mutant plants we could not detect TRKIN by immunostaining, whereas K10L2 *Trkin*(+) showed strong TRKIN staining (Figure 6). However, we did not observe TRKIN localization or TR-1 neocentromeres in plants of the Ab10 *trkin1(-) Trkin2(+)* genotype, which likely reflects the fact that *Trkin2* is expressed at very low levels (Supplementary Figure 8).

**Figure 6:**
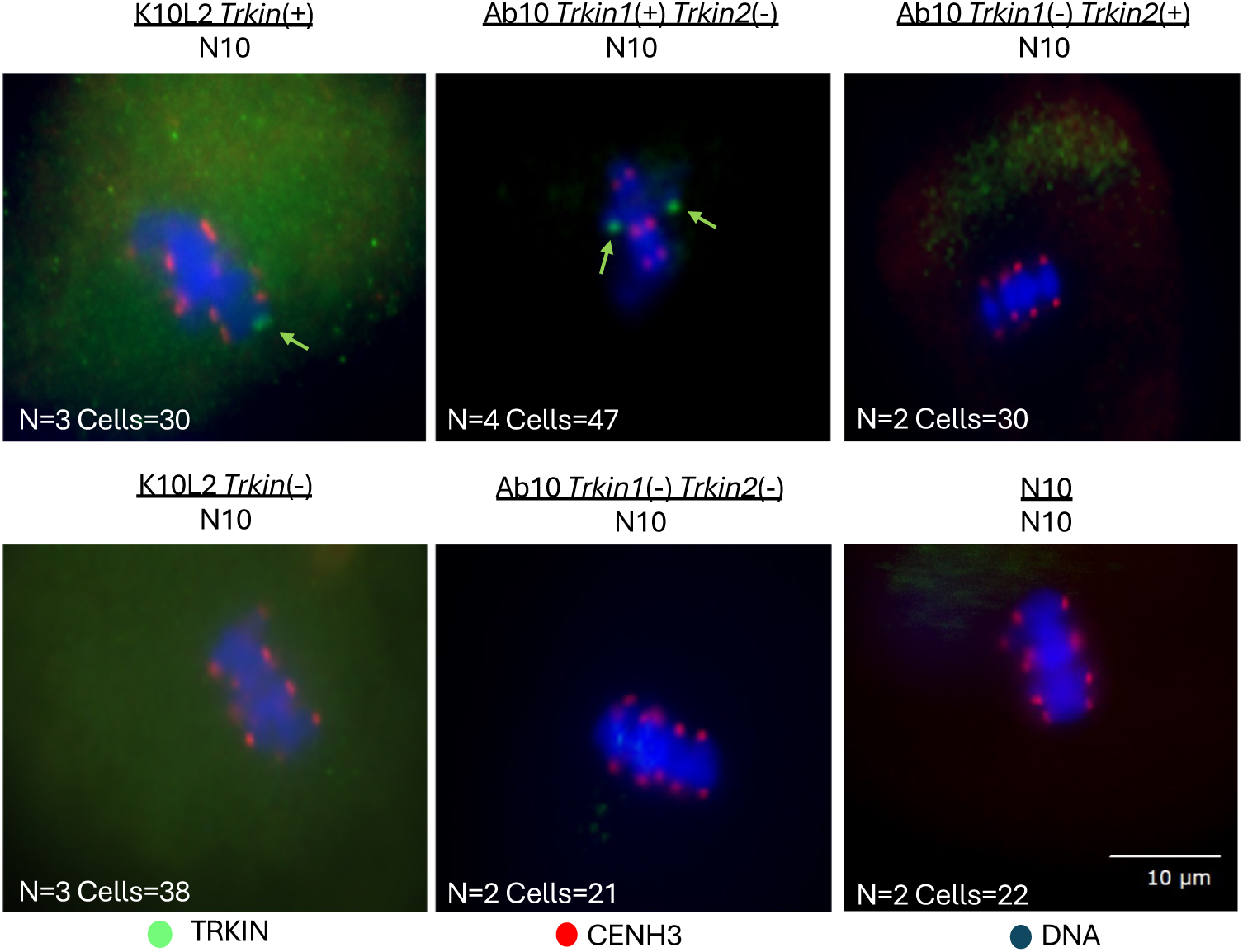
TRKIN Immunofluorescence In Various *trkin* Genotype Male Meiocytes. All images show metaphase I except for the Ab10 *trkin1*(+) *trkin2*(-) which represents metaphase II. N indicates the number of individual plants observed, cells indicate the number of appropriately staged same phenotype cells observed. CENH3 is in red, TRKIN in green, and DNA in blue. Green arrows show TRKIN staining.

**Figure 7:**
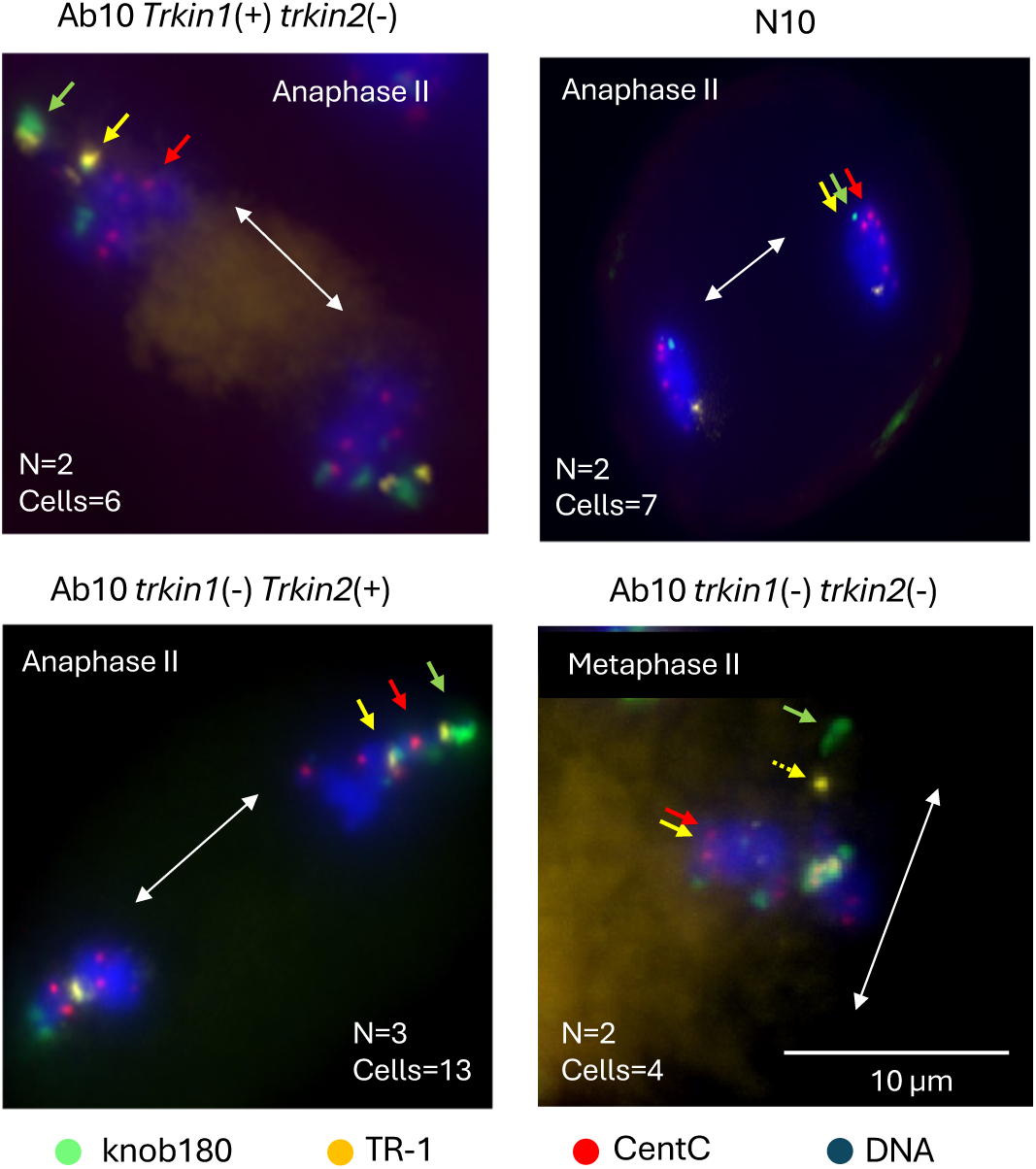
FISH for Neocentromere Activity in Various *trkin* Genotypes Male Meiocytes. All plants were homozygous for their respective genotype. All images represent male meiotic anaphase II except the Ab10 *trkin1(-) trkin2(-)* which represents male meiotic metaphase II. TR-1 and knob180 neocentromeres are known to appear in these stages (Dawe 2022). Red marks CentC, green marks knob 180, yellow marks TR-1, blue marks DNA. The white double-sided arrows indicate the spindle axis, showing which way the chromosomes were moving at the time of fixation. In the absence of TRKIN activity, TR-1 (small yellow arrows) should be located behind the centromeres (small red arrows). The yellow dot that is off the metaphase plate in the lower right panel (dotted yellow arrow) is being pulled by the large knob180 knob (this is likely Ab10 itself). N indicates the number of individual plants observed, cells indicates the number of appropriately staged same phenotype cells observed.

### The *Trkin* gene is required for K10L2 to suppress meiotic drive of Ab10

Prior work had established that when Ab10 is paired with K10L2, meiotic drive is strongly suppressed (Kanizay *et al*. 2013a). We hypothesized that K10L2 *Trkin* may be responsible for this phenomenon. Using Ab10 *Trkin1(+) trkin2(-)* and K10L2 *Trkin*(+) as positive controls, we tested the effect of *Trkin* on Ab10 and K10L2 competition. We found that when *trkin* was completely knocked out on both Ab10 and K10L2, drive was fully restored to Ab10/N10 levels (Figure 8, Supplementary Figure 9). This demonstrates that *Trkin* is necessary for K10L2 to compete with Ab10. Using reciprocal crosses, we further determined that one copy of Ab10 *Trkin1* or K10L2 *Trkin* is sufficient to fully suppress drive.

**Figure 8:**
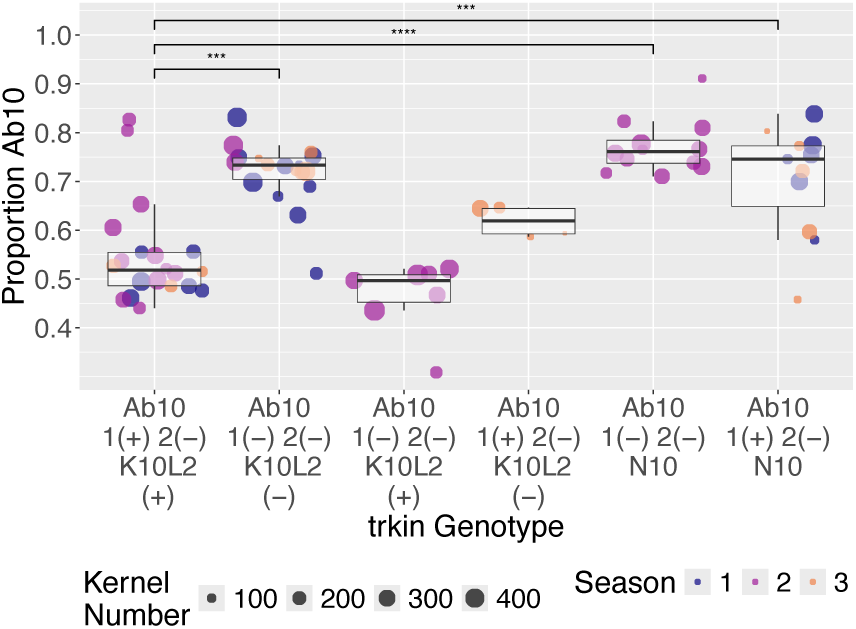
Effect of *Trkin* on the meiotic drive of Ab10 when paired with K10L2. The plot shows meiotic drive as measured by the percentage of kernels carrying the *R1* allele linked to Ab10. All plants were grown in the greenhouse in Athens, GA. Each dot represents an individual plant. Season refers to a group of plants grown at the same time. Seasons 1 and 2 were conducted in the same background while Season 3 was conducted in a different background. Season 1 and 2 of the Ab10 *trkin1(-) trkin2(-)* and *K10L2 trkin(-)* had *cas9* segregating, refer to Supplementary Figure 9 for details. The multi-way ANOVA model was Proportion Ab10 ∼ *Cas9* genotype + season + *trkin* genotype. *Cas9* genotype = F(1,63)=9.656, p=0.00; Season = F(2,63)=0.520 p=0.59726; *trkin* genotype= F(5,63)=19.495, p= 1.11e-11. Tukey’s HSD Test for multiple comparisons found that the mean value of Ab10 *Trkin1(+) trkin2(-)* / K10L2 *Trkin(+)* was significantly different from Ab10 *trkin1(-) trkin2(-)* / K10L2 *trkin(-)* (p=7.364643e-04, 95% C.I.=[3.836241-20.005130), Ab10 *trkin1(-) trkin2(-)* / N10 (p=9.996369e-09, 95% C.I.=[13.392222-31.564484) and Ab10 *Trkin1(+) trkin2(-)* / N10 (p=1.775886e-04, 95% C.I.=[5.723491-24.404879]). Only significant relationships to Ab10 *Trkin1(+) trkin2(-)* / K10L2 *Trkin(+)* are shown, refer to Supplementary Figure 9 for all significant relationships. *=<0.05, **=<0.01, ***, <0.001, ***=0.

These data suggest that Ab10 encodes its own context dependent suppressor. Ab10 with active *Trkin1* should lose most of its drive whenever it encounters K10L2, variants of K10L2 that lack *Trkin*, or any other chromosome 10 with a large TR-1 knob.

### Field and greenhouse experiments reveal no positive fitness effect of *Trkin*

Given the persistence of *Trkin* on the Ab10 haplotype, it seemed possible that it provides some benefit either through increased drive or reduced fitness effects (Buckler *et al*. 1999; Swentowsky *et al*. 2020). We tested this hypothesis by crossing our Ab10 *trkin* mutant lines as heterozygotes (*R1*-Ab10 (edited *trkin* alleles)/r1-N10) with pollen from *r1*/*r1* homozygous plants in a large, randomized field design. Drive was measured by counting kernels carrying the dominant *R1* allele, which makes the kernels purple (*r1*/*r1* is colorless). We found that Ab10 *trkin1(-) trkin2(-)* had significantly higher drive than both Ab10 single *trkin* mutants with a mean difference of 0.41% (1 - 2 +) and 0.96% (1+ 2-) (Figure 9a). These effect sizes are quite small and right at the edge of what our experiment had power to detect. We had 51.8% power to detect a 1% change in drive and 82.8% power to detect a 1.2% change in drive. These data indicate that *Trkin* is not increasing Ab10 drive under the tested experimental conditions. Instead, *Trkin* appears to decrease drive.

**Figure 9:**
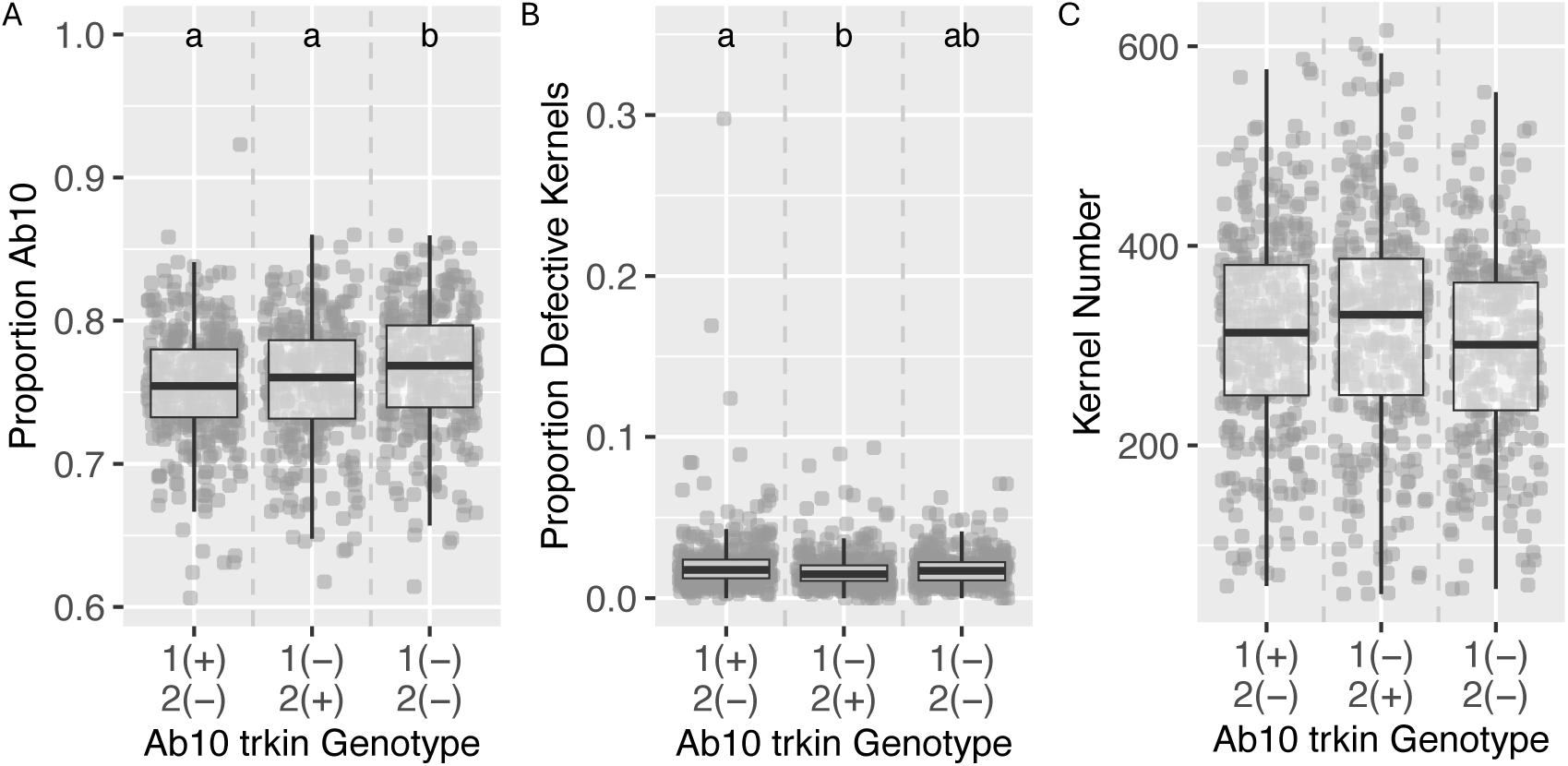
Ab10 Drive and Plant Fitness Effects of *Trkin* In Ab10 Heterozygotes. The plot shows meiotic drive as measured by the percentage of kernels carrying the *R1* allele linked to Ab10. Plants were grown in randomized order in a field in Molokai Hawaii. Each dot represents an individual plant. A. Drive: Multi-way ANOVA model was sqr (Proportion Ab10-I) ∼ field x + field y + field edge + kernel sorter + *trkin* genotype. Field x = F(1,941)=0.331, p=0.56; field y = F(1,941)=0.135, p=0.71; field edge = F(1,941)=5.475, p=0.02; kernel sorter= F(4, 941)=1.392, p=0.23; *trkin* genotype= F(2,941)=6.986, p= 0.00. B. Proportion of Defective kernels as a proxy for kernel abortion. Defective Kernels: Kruskal Wallis test model was: Proportion Defective Kernels ∼ *trkin* Genotype. H(2)=10.642, p=0.00. Wilcoxon rank sum test found A10 *Trkin1(+) trkin2(-)* (mean 0.214) was significantly different from Ab10 *trkin1(-) Trkin2(+)* (mean = 0.0173, p=0.0036), but was not significantly different from Ab10 *trkin1(-) trkin2(-)* (mean= 0.0181, p=0.1781). C. Kernel Number: Multi-way ANOVA model was Kernel Number ∼ field x + field y + field edge + kernel sorter + *trkin* genotype. Field x = F(1,941)=,1.785 p=0.18; field y = F(1,941)=3.538, p=0.06; field edge = F(1,941)=12.734, p=0.00; kernel sorter= F(4,941)=2.188, p=0.07; *trkin* genotype= F(2,941)=1.726, p=0.18.

It has previously been suggested that *Trkin* may improve Ab10 fitness by preventing anaphase segregation errors that might occur when centromeres and neocentromeres move in opposite directions on the spindle (Swentowsky *et al*. 2020). Such errors would be expected to cause increased numbers of aborted kernels. On the same ears used for testing drive, we found that Ab10 *Trkin1(+) trkin2(-)* had a significantly higher proportion of defective kernels than Ab10 *trkin1(-) Trkin2(+)* with a mean difference of 0.41%. However, Ab10 *trkin1(-) trkin2(-)* did not have a significantly different proportion of defective kernels than either single mutant (Figure 9b). We had 13% power to detect a 0.4% change and 78.2% power to detect a 0.8% change in kernel abortion. We also tested the effect of *Trkin* on the total number of kernels and found no significant differences between any genotypes (Figure 9). We had 80% power to detect down to a 30 kernel (∼8.54%) difference. These data indicate that *Trkin1* does not reduce kernel abortion or alter total kernel count.

It is well understood that Ab10 causes severe reductions in kernel count and weight when homozygous (Higgins *et al*. 2018). We hypothesized that *trkin* may be ameliorating some of the deleterious fitness effects when Ab10 is homozygous. We created an F2 population segregating for Ab10 *Trkin1(+) trkin2(-)* and Ab10 *trkin1(-) trkin2(-)* and conducted greenhouse fitness experiments. We found no significant effects on plant height, average kernel weight, or competitiveness between Ab10 haplotypes (Intra-Ab10 competition) with respect to *trkin* genotype (Supplementary Figure 10). We had 80% power to detect differences of the following magnitudes: Height = 52 cm (32% change), average kernel weight = 0.07 g (48% change), intra-Ab10 competition = 21% change. Although in this small study we only could have detected large changes, the data indicate that *Trkin1* does not improve the fitness of Ab10 in the homozygous state.

### The *Trkin1* gene does not reduce the frequency of meiotic errors in male meiosis

To test the effects of Ab10 *Trkin* on the accuracy of male meiosis, we screened Ab10 homozygous male meiocytes under the microscope for meiotic errors. Prior data demonstrated that homozygous Ab10 plants have reduced pollen viability (Higgins *et al*. 2018). We found no differences in the meiotic errors between Ab10 *Trkin1(+) trkin2(-),* Ab10 *trkin1(-) Trkin2(+)*, Ab10 *trkin1(-) trkin2(-)* lines or N10 lines (Supplementary Figure 11). We had 80% power to detect down to the following differences: Tetrad Micronuceli = 5%, Tetrad Microcyte = >0%, Dyad Micronuclei = 36%, Total Meiotic Errors = 6%. These data provide further evidence that Ab10 *Trkin1* does not reduce the frequency of meiotic segregation errors that might occur when centromeres and neocentromeres move in opposite directions on the spindle (Swentowsky *et al*. 2020).

### The *Trkin1* gene does not affect the degree of meiotic drive at an unlinked mixed knob

*Trkin* is known to activate neocentromeres throughout the genome (Dawe 2022). It seemed possible that *Trkin* behaved differently with other TR-1 knobs in the genome. To test the effect of *Trkin* on knobs elsewhere in the genome, we looked at its effect on the transmission of a large mixed knob on chromosome 4L marked by a GFP-encoding insertion that expresses in kernel endosperm (Li *et al*. 2013). We found no significant difference in segregation of the 4L knob between Ab10 with functional *Trkin1* or without functional *trkin*. We also found no difference in K10L2 *Trkin*(+) or *trkin*(-). We had 80% power to detect down to an 8% difference in segregation (Supplementary Figure 12). Together these data indicate that *Trkin* does not have an outsized effect on knobs elsewhere in the genome, just as it has little or no effect on Ab10.

### Ab10 *Trkin*(+) should not persist in maize populations and will quickly get replaced by Ab10 *trkin*(-)

The above evidence indicates that *Trkin* has a negative effect on Ab10 fitness. While it remains possible that two copies of *Trkin* have different effects or that *Trkin* has some benefit we were unable to detect, we wanted to examine the population dynamics of *Trkin* in the long-term using a modeling approach. We built on the prior Ab10 meiotic drive model (Hall and Dawe 2018) to include Ab10 *Trkin*(+), Ab10 *trkin*(-), K10L2, and N10, and examined Ab10 *Trkin*(+) dynamics in populations. Specifically, we asked three questions for a subset of parameters representative of the empirical system: (1) When and how often does Ab10 *Trkin*(+) outcompete Ab10 *trkin*(-) in a population, (2) Is the persistence of Ab10 *Trkin*(+) dominated by natural selection or genetic drift, and (3) How long does it take for Ab10 *trkin*(-) to eventually replace Ab10 *Trkin*(+) in a population?

We began with simulations following a deterministic model (assuming discrete non-overlapping generations, diploid organisms, and a single panmictic population of infinite size). We found that Ab10 *Trkin*(+) cannot invade a population at equilibrium with Ab10 *trkin*(-). Additionally, we found that Ab10 *trkin*(-) can always invade a population at equilibrium with Ab10 *Trkin*(+). Thus, unless the Ab10 *Trkin*(+) allele has some hidden or context-dependent benefit, it should not invade or segregate in a population assuming a deterministic model.

Next, we considered the strength of selection against Ab10 *Trkin*(+) reasoning that if selection is weak enough, genetic drift might dominate over natural selection in small populations. If so, genetic drift might explain the persistence of Ab10 *Trkin*(+). We calculated the selection coefficient against Ab10 *Trkin*(+) compared to Ab10 *trkin*(-) for various values of reduction in drive due to *Trkin*. Selection predominates drift if 2\**Ne*\**s* > 1, where *s* is the selection coefficient and *Ne* is the effective population size (Hartl and Clark 2007). So, we calculated 2*Ne*s for a range of reductions of drive and effective population sizes. There are almost no combinations of parameters where selection against Ab10 *Trkin*(+) would be dominated by genetic drift (2\**Ne*\**s*<1). In fact, the population size would need to be less than 100 and the reduction in drive close to zero for genetic drift dynamics to dominate: neither of which are realistic. Therefore we concluded that selection against Ab10 *Trkin*(+) is strong enough that drift cannot explain its persistence.

Though genetic drift is unlikely to prevent Ab10 *trkin*(-) from overtaking Ab10 *Trkin*(+) in a population, drift may influence how long the process takes. Given that we know both Ab10 *trkin(*-) and *Trkin(*+) segregated in wild ancestors, this suggests both have persisted for at least 8700 generations (Piperno *et al*. 2009; Swentowsky *et al*. 2020). Therefore, we assessed whether, given estimated parameters, the Ab10 *trkin(*-) might still be in the process of replacing Ab10 *Trkin(*+). Thus, we extended our deterministic model to a stochastic model (choosing genotypes from a multinomial distribution to simulate genetic drift). We asked how long it takes for Ab10 *trkin*(-) to replace Ab10 *Trkin*(+) when Ab10 *Trkin*(+) starts at a frequency of 6% (based on (Kato 1976; Kanizay *et al*. 2013a)), and Ab10 *trkin*(-) starts as a single copy. Ab10 *trkin*(-) introduced as a single copy would often be lost due to drift in a stochastic model (Haldane 1927). Figure 10a shows that the more the Ab10 *Trkin*(+) allele reduces drive, the more likely the Ab10 *trkin*(-) is to escape stochastic loss and replace Ab10 *Trkin*(+). However, in actual populations Ab10 *trkin*(-) exists so it must have escaped stochastic loss at some point (Swentowsky *et al*. 2020). Figure 10b shows the distribution for time to loss of Ab10 *Trkin*(+), given a rare Ab10 *trkin*(-) allele introduced in an Ab10 *Trkin*(+) population at equilibrium for Ab10 *Trkin*(+), K10L2, and N10 where Ab10 *trkin*(-) escaped stochastic loss. The mean time for loss of Ab10 *Trkin*(+), or the time it takes for Ab10 *trkin*(-) to replace Ab10 *Trkin*(+), is less than 500 generations. This is true if the reduction in drive is more than ∼0.01 (our empirical estimates suggest the value is more like 0.1) (Figure 9a). Therefore we concluded that Ab10 *trkin*(-) should replace Ab10 *Trkin*(+) in less than 500 generations for most parameter combinations resembling the empirical system.

**Figure 10:**
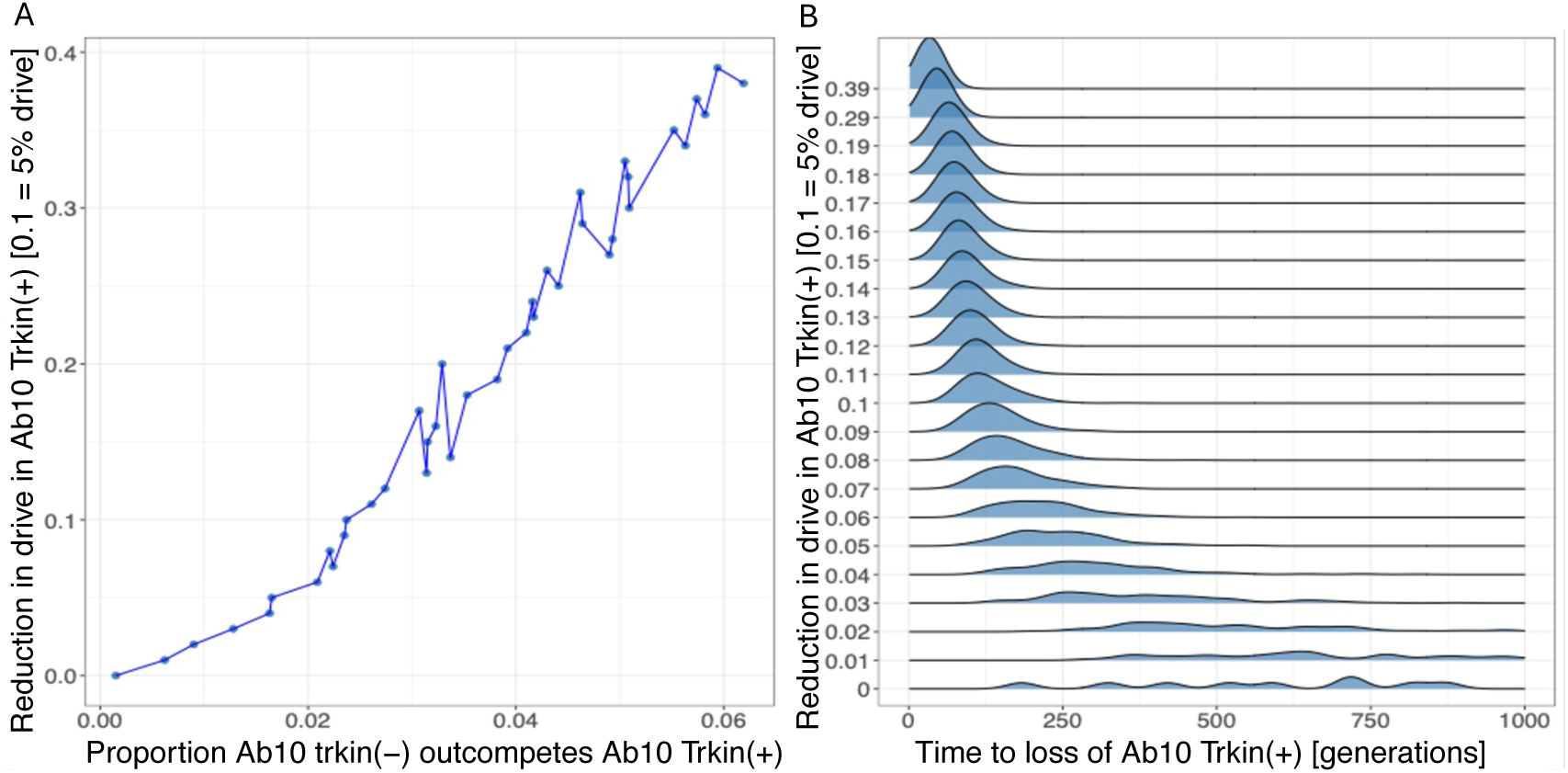
How long can Ab10 *Trkin*(+) persist in a population being invaded by Ab10 *trkin*(-)? Simulations were run stochastically, modelling drift following a multinomial distribution, at an initial frequency of 6% for Ab10 *Trkin*(+) and K10L2 and 1/Ne for Ab10 *trkin*(-) using N_e_=10,000 and for 0 < δ_1_ < 0.4. Each simulation was iterated 10,000 times. A. Proportion of realizations Ab10 *trkin*(-) successfully invades into the population and replaces Ab10 *Trkin*(+). The parameter on the y-axis is represented by δ_1_ in the model. Note that these proportions are small since Ab10 *trkin*(-) was often lost due to drift. B. Density distribution for the number of generations Ab10 *Trkin*(+) can persist in a population upon invasion by Ab10 *trkin*(-). The parameter on the Y-axis is represented by δ_1_ in the model.

The results presented above fail to explain the long-term persistence of Ab10 *Trkin*(+). They suggest that either Ab10 *trkin*(-) is very young (less than 500 generations) and is currently replacing Ab10 *Trkin*(+) or that Ab10 *Trkin*(+) confers some fitness advantage that we did not observe.

## DISCUSSION

Despite examples of *Trkin* being encoded in all three common Ab10 variants and in K10L2 (Swentowsky *et al*. 2020) and conservation of *TR-1* knobs as far as *Tripsacum*, our data provide no evidence that *Trkin* provides a selective advantage to Ab10. Instead, under the conditions we tested, Ab10 *Trkin* slightly reduces Ab10 drive and acts as an efficient suppressor of drive in the presence of K10L2. Since we only tested the function of Ab10 *Trkin1*, we cannot rule out the possibility that *Trkin1* has a positive fitness effect only in the presence of functional *Trkin2*. We can, however, confidently conclude that Ab10 *Trkin1* is sufficient to activate TR-1 neocentromeres and allow K10L2 to compete with Ab10 independently of *Trkin2*. Modeling demonstrates that, under our current understanding of the system, Ab10 *Trkin1*(+) trkin2(-) would not persist in the population if Ab10 *trkin1*(-) *trkin2*(-) were present. We propose two theories for the existence of *Trkin* on the Ab10 haplotype: an advantage either smaller than could be detected here or only apparent in untested circumstances, or that *Trkin* is in the process of being purged from the Ab10 population.

Our best estimate of *Trkin* prevalence in the Ab10 population places it at around 50% (Swentowsky *et al*. 2020). It is possible that Ab10 *trkin*(-) is a new development. Perhaps in the past, *Trkin* served a function that has been lost in the last ∼500 years and is now slowly being purged from the population. It may be that *Trkin* provides benefits to Ab10 in teosinte, but not in maize. However, maize was domesticated from teosinte ∼8700 years ago (Piperno *et al*. 2009) which our models suggest would have been ample time for *Trkin* to have been purged from the population (Figure 10b). To explain the continued presence of Ab10 *Trkin(+)* in maize it would need to be reintroduced via gene flow from teosinte, which is plausible (Yang *et al*. 2023). It is also possible that gene conversion or illegitimate recombination between Ab10 and K10L2 continuously reintroduces *Trkin* to Ab10.

K10L2 is a relatively common variant of chromosome 10 (Kato 1976; Kanizay *et al*. 2013a) and is known to function as a suppressor of Ab10 drive (Kanizay *et al*. 2013a). Our data demonstrate that the *Trkin* gene is specifically responsible for the ability of K10L2 to suppress Ab10 drive. The evolution of a suppressor on the disadvantaged allele is common in drive systems (Price *et al*. 2020). However, it is unusual and apparently paradoxical (as far as we know this is the first example) for a driving haplotype to encode its own, albeit context dependent, suppressor. The Ab10 and K10L2 drive systems are clearly complex and have had a major impact on the evolution of maize. Our data suggest that we do not yet understand the full range of contexts where Ab10 either has historically functioned or is currently functioning as a meiotic driver. Further studies of Ab10 and other chromosome 10 variants in teosinte may help provide new leads, and help us better understand the functions of *Trkin* in natural Ab10 populations.

## METHODS

### Assembly of K10L2

CI66 (PI 587148) seed was ordered from the Germplasm Resources Information Network in Ames, Iowa, and grown in the UGA Botany greenhouse in Athens, GA. Leaf tissue was sent to the Arizona Genomics Institute for DNA extraction using a CTAB method (Doyle and Doyle 1987). The sequencing library was constructed using SMRTbell Express Template Prep kit 3.0. The final library was size selected on a Blue Pippin (Sage Science) with 10-25 kb size selection. Sequencing was performed on a PacBio Revio system in CCS mode for 30 hours. We filtered reads to a quality of 0.99 or greater and converted them to fastq format using bamtools v2.5.2 and bedtools 2.30.0 respectively (Quinlan and Hall 2010; Barnett *et al*. 2011). We ran hifiasm v0.19.6 with post joining disabled to assemble the raw reads into contigs (Cheng *et al*. 2021). We identified the K10L2 haplotype by using BLAST v 2.13.0 to identify the contig with homology to the *Trkin* cDNA sequence (Swentowsky *et al*. 2020). Using BLAST v 2.13.0 we determined that the contig bearing *Trkin* also contained two large TR-1 knobs. Using the integrated genome viewers (IGV) motif finder we determined that the *Trkin* bearing contig ended in 7,674 bp of telomere sequence indicating it was fully assembled (Thorvaldsdóttir *et al*. 2013). The *Trkin* bearing contig had no homology to the *colored1* gene, which marks the beginning of the Ab10 haplotype. To ensure all the chromosome 10 haplotypes were comparable we chose to manually merge the *colored1* gene bearing contig with the contig containing the otherwise complete K10L2 haplotype. Using BLAST v 2.13.0 we identified the contig bearing the colored1 gene (B73 v5 Zm00001eb429330) and merged it to the *trkin* bearing contig with an interceding 100N gap using RagTag v2.1.0 (Alonge *et al*. 2022). All other contigs were left unaltered.

### Assembly of B73-Ab10 v2

We chose to generate a new Ab10 assembly as there had been significant methodological advances since the generation of the first assembly (Liu *et al*. 2020). We used the same high molecular weight genomic DNA that was used in the B73-Ab10 v1 assembly (Liu *et al*. 2020). The sequencing library was constructed using SMRTbell Express Template Prep kit 2.0. The sequencing library was prepared for sequencing with the PacBio Sequel II Sequencing kit 2.0 for HiFi libraries and sequenced in CCS mode at the UGA Georgia Genomics and Bioinformatics Core facility. This data was integrated into the previously published assembly pipeline to produce the v2 assembly (Liu *et al*. 2020).

### Comparison of the B73-Ab10 v1 and B73-Ab10 v2 Haplotypes

B73-Ab10 v1 and B73-Ab10 v2 were compared using Mummer v4.0.0 with a minimum length (-m) of 300 and computed all matches not only unique ones (--maxmatch) (Marçais *et al*. 2018; Liu *et al*. 2020). Plots were generated using R v4.3.1.

### Annotation of Ab10 and K10L2

The assemblies described above were annotated for repeats and masked using RepeatMasker v4.1.5 in conjunction with the maize repeat library (Smit AFA., Hubley R., Green P. 2015; Ou 2020). All available short read mRNA sequencing data was downloaded for Ab10 (Liu *et al*. 2020) and K10L2 (Swentowsky *et al*. 2020) respectively. Reads were trimmed with Trimmomatic v0.39 (Bolger *et al*. 2014). These reads were then aligned to their respective genomes using HiSat2 v3n-20201216 (Kim *et al*. 2019). The resulting files were converted to a bam format and sorted using samtools v1.17 (Kim *et al*. 2019; Danecek *et al*. 2021). These alignments were used as expression evidence and the Viridiplantae partition of OrthoDB was used as protein evidence in an annotation using BRAKER v3.0.8 (Kuznetsov *et al*. 2023; Gabriel *et al*. 2024). Trinity v2.15.1 and StringTie v2.2.1 were used to assemble a de novo and reference guided transcriptome from the compiled RNAseq data for Ab10 and K10L2 respectively (Haas *et al*. 2013; Pertea *et al*. 2015). These transcriptomes were combined and converted to a comprehensive transcriptome database using PASA v2.5.3 (Haas *et al*. 2003). The resulting comprehensive transcriptome database was used to polish and add UTRs to the BRAKER derived gene annotation file in three rounds of PASA v2.5.3 (Haas *et al*. 2003). We found that the *Trkin* bearing region on Ab10 and K10L2 has an average percent identity of 98.5% for aligned regions (Figure 2). However, the annotated genes were quite different. In order to improve the annotations we used Liftoff v1.6.3 to reciprocally update the annotations in the *Trkin* bearing region on both haplotypes (Shumate and Salzberg 2021). We then extracted only genes that were included in the liftoff annotation using bedtools v2.31.0 and incorporated them (Quinlan and Hall 2010). Genes added in this way have names starting with gA in the K10L2 annotation and gK in the Ab10 annotation. We extracted the CDS and cDNA sequences for both haplotypes using AGAT v1.1.0 (Dainat 2020) Finally, we extracted and functionally annotated the final protein sets using EnTAP v1.0.0 with the nr, Refseq, and Uniprot databases (O’Leary *et al*. 2016; Hart *et al*. 2020; Sayers *et al*. 2022; UniProt Consortium 2023).

### Determination of Ab10 knob180 Knob Size

We obtained illumina sequence reads for terminal deletions of Ab10 in the W23 inbred background that either did or did not contain the large knob180 knob on the distal most end (Brady *et al*. 2024). We quantified knob180 repeat abundance in raw illumina short reads as described in (Hufford *et al*. 2021). In brief, we used seqtk v 1.2 to convert the read files to fasta format, used BLAST v2.2.26 to identify reads with homology to knob180, and bedtools merge v2.30.0 to combine overlapping hits (Quinlan and Hall 2010; Camacho *et al*. 2023; “seqtk” 2023). Using a custom R script, we filtered to hits 30 bp or longer, summed the lengths of all hits and divided that value by the average coverage of the library to obtain the Mb value of knob180 in each library. We then subtracted the value of the intact W23-Ab10 from the sample which did not contain the large knob180 knob to obtain the estimated size of the knob180 knob on Ab10. We repeated this process for TR1 and CentC as negative controls.

### Comparison of Sequence Homology Between Ab10 and K10L2

All possible pairwise comparisons of chromosome 10 haplotypes were made using Mummer v4.0.0 with a minimum length (-m) of 300 and computed all matches, not only unique ones (--maxmatch). Self by self comparisons were run using the --nosimplify flag (Marçais *et al*. 2018). Plots were generated using R v4.3.1.

To assess the completeness of the nrpd2/e2 gene homologs we extracted all annotated copies coding sequence using AGAT v1.1.0 (Dainat 2020). We then aligned all copies to the nrpd2/e2 coding sequence from the B73v5 assembly using Geneious Prime v 2022.0.2 geneious algorithm (“Geneious 2022.0.2” 2022) (Zm00001eb068960) (Hufford *et al*. 2021). We identified functional domains in the nrpd2/e2 coding sequence using NCBI conserved domain search (Wang *et al*. 2023).

### Comparison of trkin CDS

The newly annotated *Trkin* gene was identified by overlap with the BLAST v 2.13.0 hits for *Trkin* cDNA (Swentowsky *et al*. 2020) against the newly assembled references (Camacho *et al*. 2023). The associated CDS was extracted from the CDS file for the respective genomes produced using AGAT v1.1.0 (Dainat 2020). The CDS sequences were aligned using Geneious Prime v 2022.0.2 geneious algorithm (“Geneious 2022.0.2” 2022). Protein domain locations were determined using NCBI conserved domain search, the cNLS mapper, and the MPI Bioinformatics toolkit (Kosugi *et al*. 2009; Gabler *et al*. 2020; Wang *et al*. 2023).

To better understand the relationship between the *Trkin* alleles we chose to make a phylogenetic tree using the protein motor domain. Unfortunately, TRKIN does not share sufficient homology with similar proteins to use its entire length. (Swentowsky *et al*. 2020). We used NCBI conserved domain search (Wang *et al*. 2023) to identify the motor domain in all the *Trkin* alleles as well as *Drosophila melanogaster* Ncd (Uniprot P20480) and *Zea mays Dv1* (B73 v5 annotation Zm00001eb069600). We selected *Zea mays Dv1* as it is the most closely related gene to *Trkin* (Swentowsky *et al*. 2020). We selected *Drosophila melanogaster* Ncd to act as an outgroup. We used geneious prime v2022.0.2 to (“Geneious 2022.0.2” 2022) perform a MUSCLE alignment of all 4 motor domains and used the geneious tree builder to create a Neighbor-Joining tree using the Jukes-Cantor model. We set *Ncd* as the outgroup and performed 10000 bootstrap replicates. Numbers at nodes indicate the percent of replicate trees supporting that node.

### Comparison of Gene Orthologs

Gene orthology between the three variants of the chromosome 10 haplotype was compared as described in (Brady *et al*. 2024). For the purposes of this analysis, the beginning of each haplotype was determined to be the location of the *colored1* gene. Plots were generated using R v4.3.1.

### GO term enrichment analysis

We isolated the non-shared region, defined as those areas with no consistent synteny or homology to N10 as determined by the gene ortholog analysis and sequence comparisons, for both Ab10 and K10L2. These genes were tested against the remaining portions of the genome for GO term enrichment using topGO (Adrian Alexa 2024). The Ab10 non-shared region contains several known duplicated genes that heavily influence the results. All known arrayed gene duplicates were collapsed down to a single copy. The two copies of *Trkin* were both included.

### Expression of Trkin

We obtained RNA sequencing data for Ab10 and K10L2 from (Swentowsky *et al*. 2020). We trimmed reads using Trimmomatic v0.39 (Bolger *et al*. 2014) and aligned them to the Ab10 v1 reference (Liu *et al*. 2020) using HiSat2 (Kim *et al*. 2019) and processed the output using Samtools v1.9 (Danecek *et al*. 2021). We used the R package featureCounts to determine the expression for each annotated gene (Liao *et al*. 2014). We then calculated the transcripts per million (TPM) for Ab10 *Trkin1* and Ab10 *Trkin2* in all samples requiring a mapping quality of 20. We summed the TPM of Ab10 *trkin1* and *trkin2* for easy comparison between Ab10 and K10L2.

To assess the expression of Ab10 *Trkin1* and *Trkin2* separately we assessed expression at the individual exon level. We obtained RNA sequencing data for 10 tissues of the B73-Ab10 inbred (Liu *et al*. 2020). We aligned them to the Ab10 v2 reference generated here using HiSat2 (Kim *et al*. 2019). We filtered the alignments to a mapping quality of 20 and required no mismatches. We then used the R package featureCounts to determine the expression of each annotated exon (Liao *et al*. 2014). We then calculated the TPM for only the *Trkin* exons containing SNPs (7 and 8) in all samples (Figure 3). We used a Welch two sample t-test to determine statistical significance between the two alleles.

### Construction and transformation of a plasmid expressing Cas9 and guide RNAs

A CRISPR plasmid expressing Cas9 and three guide RNAs targeting *trkin* was constructed using a pTF101.1 binary plasmid (Paz *et al*. 2004) with similar components as previously used for gene editing in maize (Wang *et al*. 2021). In particular, it utilizes 1991 bp of a maize polyubiquitin promoter and UTR region (GenBank, S94464.1) to drive expression of Cas9 from *Streptococcus pyogenes* flanked by an N-terminal SV40 NLS and a C-terminal VirD2 NLS and followed by a polyadenylation signal provided by a *nopaline synthase* (*NOS)* terminator sequence from *Agrobacterium tumefaciens*. The Cas9 DNA sequence was codon optimized for maize as described previously except that it did not include the potato ST-LS1 intron (Svitashev *et al*. 2015). The three guide RNAs were transcribed by three individual U6 promoters from maize and rice with two gRNAs targeting *Trkin* exon 3 (GTCTGGAGGCCAATGAGCACG and GAAAGCTTTTGCGGCCTCTGG) and one targeting exon 4 (GCCTACACAAGTAAACAGAT). These target sequences were selected using CHOPCHOP v3 (Labun *et al*. 2019). See Supplemental File 1 for complete plasmid sequence and annotations. Gene synthesis and cloning was performed by GenScript (www.genscript.com), and transformation was performed by the Iowa State University Plant Transformation Facility.

### Genotyping for trkin mutants

All genotyping DNA extractions were performed using a CTAB protocol (Clarke 2009). Polymerase chain reactions were performed using Promega GoTaq Green Master Mix (M7123). The Ab10 *trkin1* and K10L2 *trkin* edits were identified using the same primers (trkin_EX3 and trkin_EX4), Ab10 *trkin2* was detected using a separate pair of primers (Ptrkin_EX3, Ptrkin_EX4) (Supplementary Table 6). Edits were confirmed by purifying the PCR reaction via Omega Bio-Tek Mag-Bind RxnPure Plus beads (M1386-01) using a 1:1 ratio and Sanger sequencing by Eton Biosciences. The competition assay plants were genotyped using primers specific to an indel in an intron of the *Trkin* gene (K10L2) (Supplementary Table 6). All lines were checked for Cas9 using specific primers (Supplementary Table 6). All reactions were conducted with slightly different temperature profiles and concentrations detailed in Supplementary Table 6.

### Immunofluorescence and FISH

Both Immunofluorescence and FISH were performed as described in (Swentowsky *et al*. 2020).

### Competition Assay

To assess the effect of *Trkin* on the ability of K10L2 to suppress Ab10 drive we used plants in the same background that had one copy of Ab10 and one copy of K10L2 with varying *trkin* genotypes. In all cases Ab10 was marked by a dominant functional allele of *the colored 1* (*R1)* and K10L2 was marked by a recessive mutant allele (*r1*). We crossed these plants as the female to an r1/r1 male and scored segregation of the R1 allele. The background used contained the *C1* allele and was thus appropriate for tracking the *R1* allele. All experiments were conducted in in the UGA Botany greenhouse (Athens, GA) across 3 seasons. In the case of K10L2 *trkin*(-) one season of the experiment had Cas9 segregating thus making it impossible to determine what *trkin* mutation was present. These are indicated in (Supplementary Figure 9).

Results were analyzed using an ANOVA. Plots were generated using R v4.3.1.

### Assessment of Ab10 Heterozygous Drive and Fitness

To determine the effect of *Trkin* on Ab10 drive we generated plants heterozygous for Ab10 and N10 with various *trkin* genotypes in the same genetic background. Friendly Isles Growing planted all plants in Molokai Hawaii in randomized rows of 15 kernels with every other row being an *r1/r1* male. No border corn was used, but edge effects were included in the final statistical model. All Ab10 bearing plants were detasseled, and allowed to open pollinate with the *r1/r1* males. Upon completion of the growing season Friendly Isles Growing harvested all female plants and sent them to the University of Georgia for processing. All ears were scored for defective kernels, a proxy for aborted kernels, defined as clearly defective kernels surrounded by otherwise healthy kernels with no other explanation. These criteria were selected to exclude insect damage, vivipary, and kernel loss during shipment. We shelled the ears and sorted them by color (dark pigmented *R1* and yellow *r1*). The seeds in each packet were counted using an International Marketing and Design Corp. Programmable Packeting Model 900-2 seed counter with the fast set to 7.2 and the slow set to 0.

The meiotic drive data were found to violate the criteria for an ANOVA, so we square root transformed the data to improve its fit which did not fully satisfy the statistical assumptions for a linear relationship, skew, and kurtosis, but came reasonably close. We chose to proceed with the ANOVA as the residuals appeared normally distributed and alternative statistical methods didn’t offer the ability to account for the necessary number of variables. We included the following covariates in the model: field x coordinate, field y coordinate, edge of field, individual who sorted the kernels. The kernel abortion data was very far from a normal distribution so a kruskal-wallis test was used. The total kernel number data were analyzed using an ANOVA and met all assumptions. We included the following covariates in the model: field x coordinate, field y coordinate, edge of field, individual who sorted the kernels. Refer to Figure 9 for the full model used for each test.

### Assessment of Ab10 homozygous fitness

To assess the effect of *Trkin* on Ab10 fitness we created an F2 mapping population segregating for Ab10 *Trkin1(+) trkin2(-)* and Ab10 *trkin1(-) trkin2(-)*. We grew 39 F2 plants and scored them for their *trkin1* genotype. We used a chi square test to check for deviation from a Mendelian segregation pattern. Plants were placed in a randomized order and grown to maturity in the UGA Botany greenhouse. They were allowed to open-pollinate amongst themselves. We measured plant height, and average kernel weight as proxies for plant fitness. We also scored total kernel count, but the experiment was underpowered to detect an effect of any magnitude. All data was analyzed using an ANOVA. Plots were generated using R v4.3.1.

### Effect of Trkin on male meiotic errors

We scored Ab10 homozygous plants with different *trkin* genotypes for meiotic errors using the slides prepared for FISH as described above. A meiotic error was defined as a micronucleus in a dyad or tetrad, or a microcyte in a dyad or tetrad (Supplementary Figure 11). Counts of meiotic errors were normalized against the total count of same stage cells observed. Results were analyzed using an ANOVA. Plots were generated using R v4.3.1.

### Effect of Trkin on unlinked mixed knob

We ordered a line carrying a marker gene expressing GFP from a zein promoter (Li *et al*. 2013) that is closely linked to the knob on chromosome 4L (tdsgR106F01) from the Maize Genetics Cooperation Stock Center, Urbana, Illinois. We generated lines heterozygous for Ab10 or K10L2 with various *trkin* genotypes where the GFP insertion was linked to the knob and the opposite chromosome 4L was from the inbred Ms71 (PI 587137), which lacks a knob on 4L (Albert *et al*. 2010). Cas9 was segregating in the families used for these experiments so it wasn’t possible to determine the exact allele used. However, all plants were derived from an individual with a *trkin* null mutation making it extremely likely that all plants, even those carrying Cas9, carry a *trkin* null mutation as well. We then crossed these lines as the female to Ms71 and scored the resulting kernels for GFP fluorescence under visible blue light using a Dark Reader Hand Lamp and Dark Reader Glasses (Clare Chemical Research #HL34T). All data were analyzed using an ANOVA. Plots were generated using R v4.3.1.

### Modeling the effect of trkin on Ab10 population dynamics

We model the system as a single locus where four alleles (Ab10 *Trkin*(+), Ab10 *trkin*(-), K10L2 and N10) are segregating. We initially assumed finite population sizes, discrete non-overlapping generations, diploid organisms, a single panmictic population, and that all individuals have the same number of offspring. We introduced stochasticity later. We assumed the N10/N10 homozygote is the wild-type genotype and has maximal fitness. We assumed that all heterozygotes experience drive during ovule production; pollen production follows Mendelian transmission and Ab10 *Trkin*(+), Ab10 *trkin*(-) and K10L2 alleles bear a fitness cost (Table 1, Table 2). Ab10 drives against N10 (drive strength: d_1_) and K10L2 (drive strength: d_3_). K10L2 drives against N10 (drive strength: d_2_). The *Trkin*(+) allele suppresses Ab10 drive by an amount of δ_1_ (0<δ_1_<d_1_).

**Table 1.** Ab10 *trkin* Modeling. Fitness and proportion of ovules and pollen produced by each genotype.

| Genotype | Proportion ovules |  |  |  | Proportion pollen |  |  |  | Fitness |
| --- | --- | --- | --- | --- | --- | --- | --- | --- | --- |
|  | N10 | Ab10<br>*Trkin*<br>(+) | Ab10<br>*trkin*<br>(-) | K10L2 | N10 | Ab10<br>*Trkin*<br>(+) | Ab10<br>*trkin*<br>(-) | K10L2 |  |
| N10<br>N10 | 1 |  |  |  | 1 |  |  |  | 1 |
| N10<br>Ab10 *trkin*(-) | $(1-d_1)/2$ | | $(1+d_1)/2$ | | $1/2$ | | $1/2$ | | $1-h_a a$ |
| N10<br>Ab10 *Trkin*(+) | $(1-d_1+\delta_1)/2$ | $(1+d_1-\delta_1)/2$ | | | $1/2$ | $1/2$ | | | $1-h_a a$ |
| N10<br>K10L2 | $(1-d_2)/2$ | | | $(1+d_2)/2$ | $1/2$ | | | $1/2$ | $1-h_k k$ |
| Ab10 *trkin*(-)<br>Ab10 *trkin*(-) | | | 1 | | | | 1 | | $1-a$ |
| Ab10 *Trkin*(+)<br>Ab10 *trkin*(-) | | $1/2$ | $1/2$ | | | $1/2$ | $1/2$ | | $1-a$ |
| Ab10 *Trkin*(+)<br>Ab10 *Trkin*(+) | | | 1 | | | | 1 | | $1-a$ |
| K10L2<br>K10L2 | | | | 1 | | | | 1 | $1-k$ |
| K10L2<br>Ab10 *Trkin*(+) | $(1-d_3)/2$ | $(1+d_3)/2$ | | | $1/2$ | $1/2$ | | | $(1-h_a a)^*(1-h_k k)$ |
| K10L2<br>Ab10 *trkin*(-) | $(1-d_3)/2$ | | $(1+d_3)/2$ | | $1/2$ | | $1/2$ | | $(1-h_a a)^*(1-h_k k)$ |

**Table 2:** Ab10 *Trkin* Model Parameters. Parameters used in the model (All parameters range between 0-1 except δ_1_, δ_1_ ranges between 0-d_1_).

| Variable/<br>Parameter | Description |
| --- | --- |
| $d_1$ | Drive strength of Ab10 against N10 |
| $\delta_1$ | Amount of Ab10 drive suppressed by *trkin*(+) |
| $d_2$ | Drive strength of K10L2 against N10 |
| $d_3$ | Drive strength of Ab10 against K10L2 |
| $a$ | Fitness cost of Ab10 homozygote |
| $h_a$ | Dominance coefficient for Ab10/N10 |
| $k$ | Fitness cost of K10L2 homozygote |
| $h_k$ | Dominance coefficient for K10L2/N10 |

Let p_m_^+^, p_f_^+^, p_m_^-^, p_f_^-^, q_m_, and q_f_ denote the frequencies of the Ab10 *Trkin*(+), Ab10 *trkin*(-), and K10L2 alleles in pollen and ovules respectively in one generation. Then, the frequencies of the alleles in the next generation can be given by –

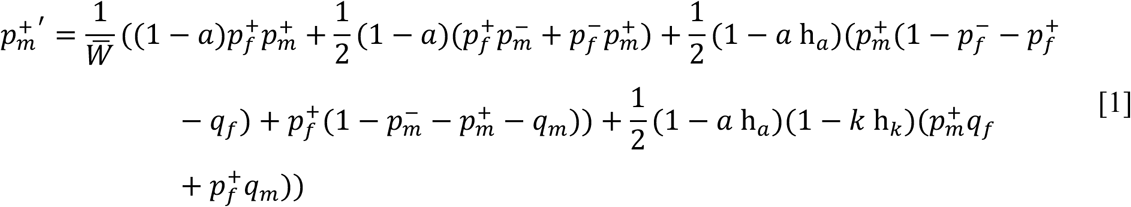

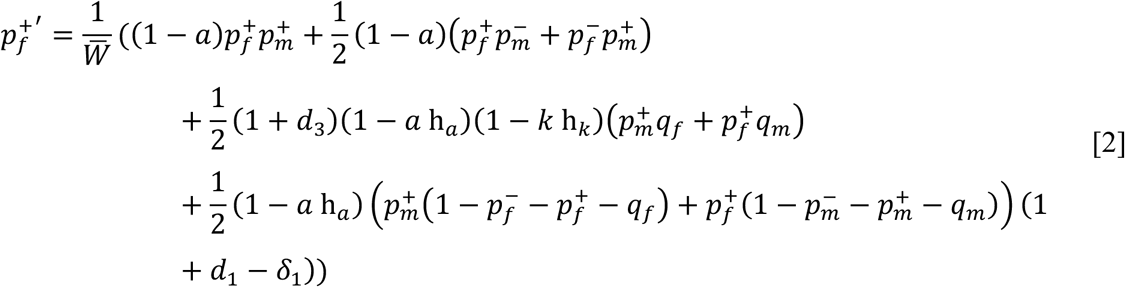

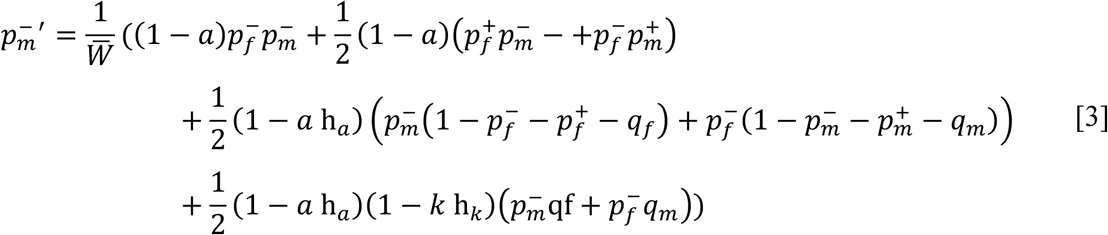

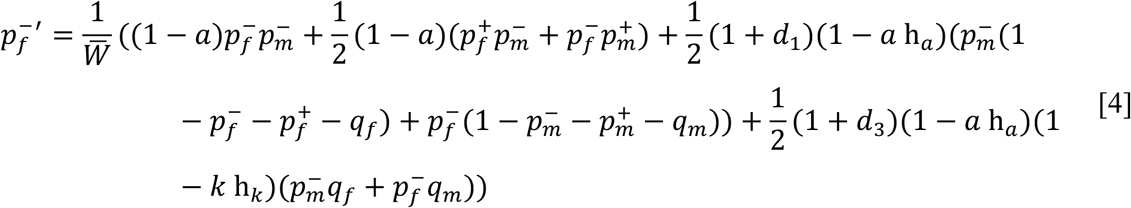

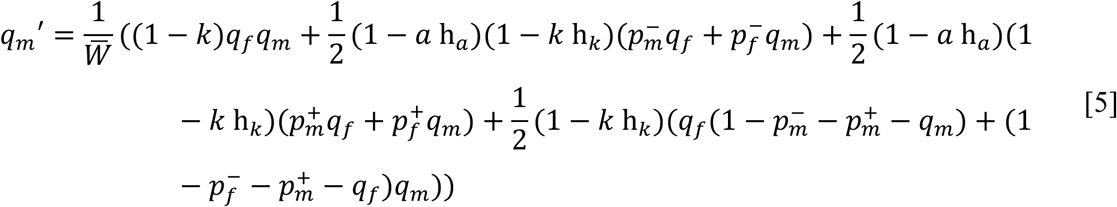

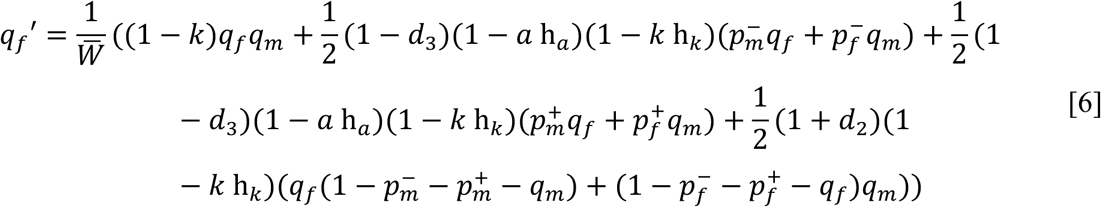

Here, the mean fitness 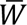 can be calculated using –

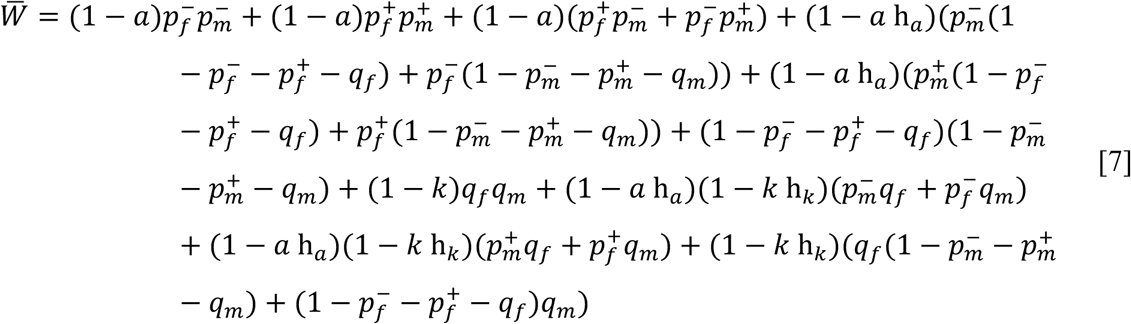

The frequency of N10 allele in pollen and ovules can be calculated using 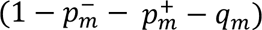 and 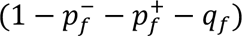 respectively. We track the frequencies separately in the two sexes such that the frequencies in males and females each add up to 1, and the population always has equal sex-ratios.

We use a subset of parameters for the simulations based on empirical observations from the maize system – h_a_ = 0.25, h_k_ = 0.2, a = 0.6, k = 0.225, d_1_ = 0.4 (drive strength of Ab10 against N10 = 70%), d_2_ = 0.1 (drive strength of K10L2 against N10 = 55%), d_3_ = 0.1 (drive strength of Ab10 against K10L2 = 55%) (Kanizay *et al*. 2013a; Higgins *et al*. 2018).

At this parameter subset, at δ_1_=0, at equilibrium, both Ab10 and K10L2 persist at a frequency of 5% each and the frequencies of Ab10 *Trkin*(+) and Ab10 *trkin*(-) are equal (deterministically).

### Testing the range of d_1_ where Ab10 Trkin(+) and Ab10 trkin(-) can invade a population

We ran these simulations deterministically for a range of δ_1_ (0 < δ_1_ < 0.4) using an effective population size, N_e_ of 10,000 (Tittes *et al*. 2021) for 5000 generations (sufficient to reach equilibrium) with initial frequencies of Ab10 *Trkin*(+) and K10L2 at 5%, and Ab10 *trkin*(-) at 1/N_e_ (equal frequencies in both sexes). At any δ_1_ > 0, Ab10 trkin(-) always invades the population and replaces Ab10 *Trkin*(+).

We also tested for the invasion of Ab10 *Trkin*(+) similarly by starting the simulations with initial frequencies of Ab10 *trkin*(-) and K10L2 at 5%, and Ab10 trkin(+) at 1/N_e_ (equal frequencies in both sexes). For any value δ_1_, Ab10 *Trkin*(+) could never invade the population.

This suggests that the selection against Ab10 *Trkin*(+) is strong to prevent its invasion in a population containing Ab10 trkin(-) and Ab10 *trkin*(-) can invade a population containing Ab10 *Trkin*(+) and replace it.

### Testing the strength of selection for a range of d_1_ and calculating the selection coefficients such that 2N_e_ s < 1 (nearly neutral zone)

For the calculation of the relative selective benefit (*s*) for Ab10 *trkin*(-), we ran the simulations for a range of δ_1_ (0 < δ_1_ < 0.4) for 5000 generations (sufficient to reach equilibrium) with initial frequencies of Ab10 *Trkin*(+) and K10L2 at 1/N_e_, and Ab10 *trkin*(-) at 0. Then, after 5000 generations, we introduced Ab10 *trkin*(-) at a frequency of 1/Ne (only in females) into the population at equilibrium. Then, we ran the simulation for one more generation and calculated the relative selective benefit of Ab10 *trkin*(-), *s* using allele frequencies after generation 5000 using –

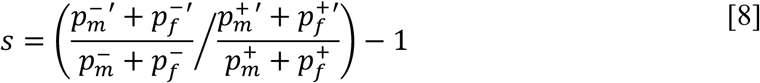

This ‘*s*’ was used to calculate the 2N_e_ *s* parameter for a range of values of N_e_ (10^2^ < Ne < 10^4^) and δ_1_ (0 < δ_1_ < 0.4). We found that 2 N_e_ *s* < 1 only for a very small subset where d_1_ < 0.01 and N_e_ ∼ 100 (The approximate value of δ_1_ from empirical observations in the maize system should be ∼ 0.1)(Figure 9a). This suggests that selection against Ab10 *Trkin*(+) is strong and it could not be maintained in the population by drift (since 2 N_e_ *s* >> 1). This would imply that Ab10 *Trkin*(+) could not persist in the population in the presence of Ab10 *trkin*(-). Ab10 *Trkin*(+) is probably older than Ab10 *trkin*(-) and could be in the process of being replaced from the populations by invasion from Ab10 *trkin*(-).

### Testing how long Ab10 Trkin(+) can persist in a population that is being invaded by Ab10 trkin(-)

We ran these simulations stochastically (modelling drift following a multinomial distribution) at N_e_=10,000 and for a range of δ_1_ (0 < δ_1_ < 0.4) (Tittes *et al*. 2021). We started our populations at an initial frequency of 6% for Ab10 *Trkin*(+) and K10L2 and 1/Ne for Ab10 *trkin*(-) (equal frequencies in both sexes). For each parameter value, each simulation was run 10,000 times, as Ab10 *trkin*(-) was often lost due to drift.

For the subset of simulations where Ab10 *trkin*(-) could successfully invade and replace Ab10 *Trkin*(+), we looked at the time taken for loss of Ab10 *Trkin*(+) from the population (Figure 10 B). For most values of δ_1_, Ab10 *Trkin*(+) was lost within 500 generations. From empirical estimates, δ_1_ ∼ 0.1, thus, Ab10 *Trkin*(+) would be expected to persist for ∼ 200 generations (Figure 9a).

We also looked at the proportion of times Ab10 *trkin*(-) (escaping stochastic loss due to drift) could successfully invade the population and outcompete Ab10 *Trkin*(+) (Figure 10 A). This proportion was small and for δ_1_ ∼ 0.1, about 2.5% of the times Ab10 *trkin*(-) could escape stochastic loss and outcompete Ab10 *Trkin*(+).

## FIGURE LEGENDS

**Supplementary Figure 1:**
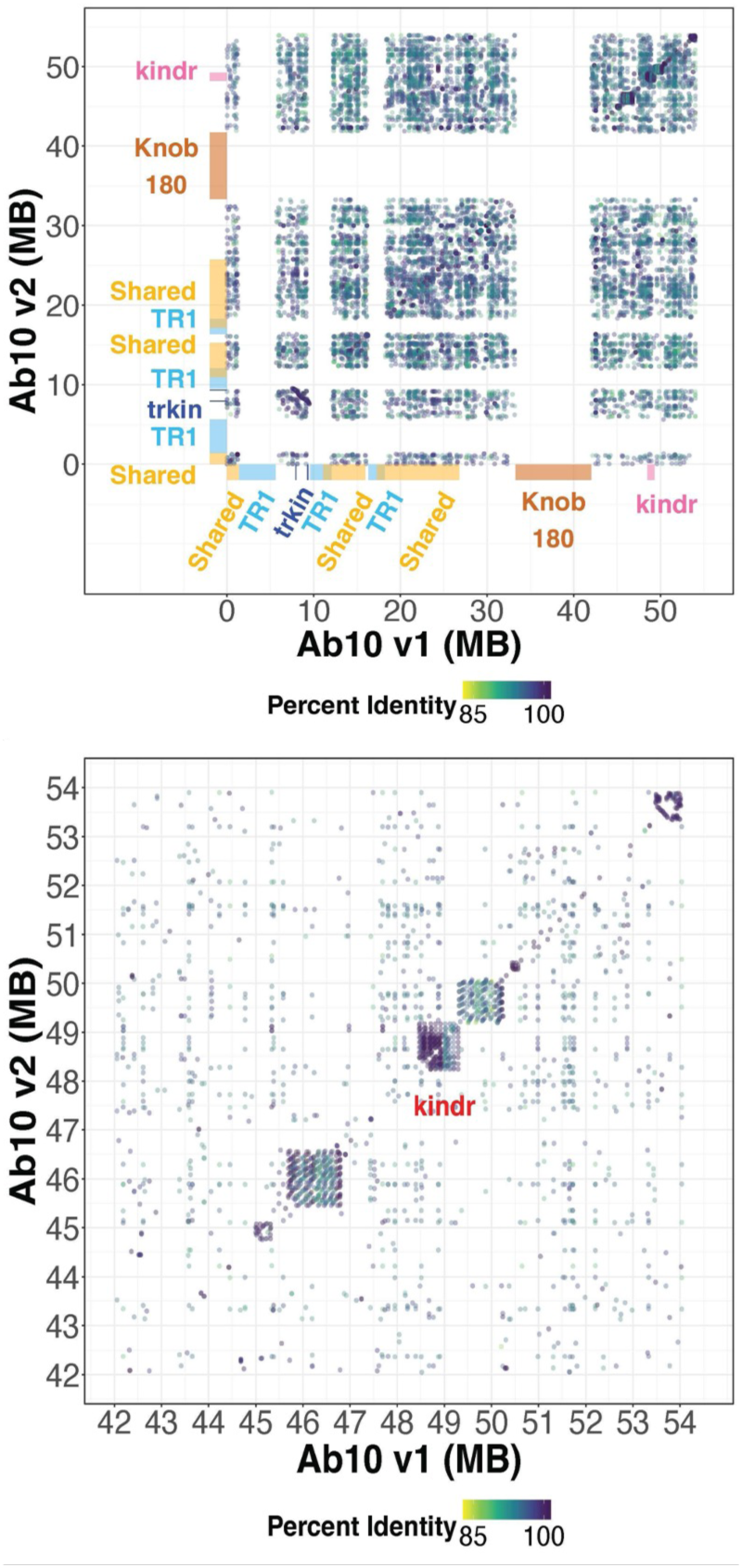
Sequence Comparison of Chromosome 10 Haplotypes. Each dot marks the start of a maximal unique match (MUM) of at least 300bp long between the B73-Ab10 v1 and B73-Ab10 v2 genomes Ab10 haplotype, which begin at the *colored1* gene (Marçais *et al*. 2018). B73-Ab10 v1 refers to the first Ab10 assembly (Liu *et al*. 2020), B73-Ab10 v2 refers to the assembly generated here. The color of each dot represents the percent identity of that match. All large knob arrays were removed for the sake of clarity. Relevant regions of each genome are marked.

**Supplementary Figure 2:**
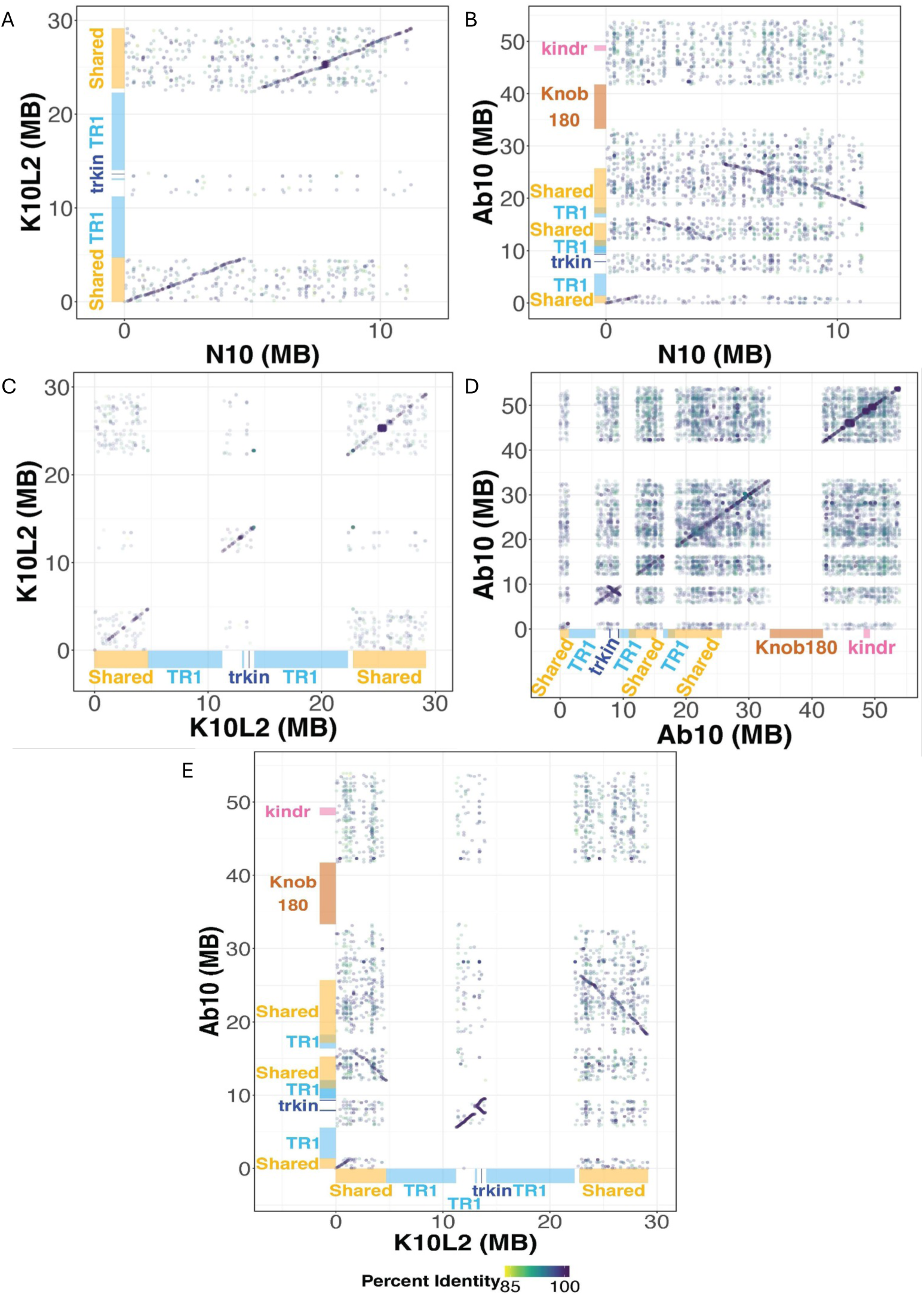
Sequence Comparison of Chromosome 10 Haplotypes. Each dot marks the start of a maximal unique match (MUM) of at least 300 bp long between various chromosome 10 haplotypes all of which begin at the *colored1* gene (Marçais *et al*. 2018). N10 refers to the B73 v5 assembly (Hufford *et al*. 2021), Ab10 and K10L2 refer to the assemblies generated in this work. The color of each dot represents the percent identity of that match. All large knob arrays were removed for the sake of clarity. Relevant regions of each haplotype are marked.

**Supplementary Figure 3:**
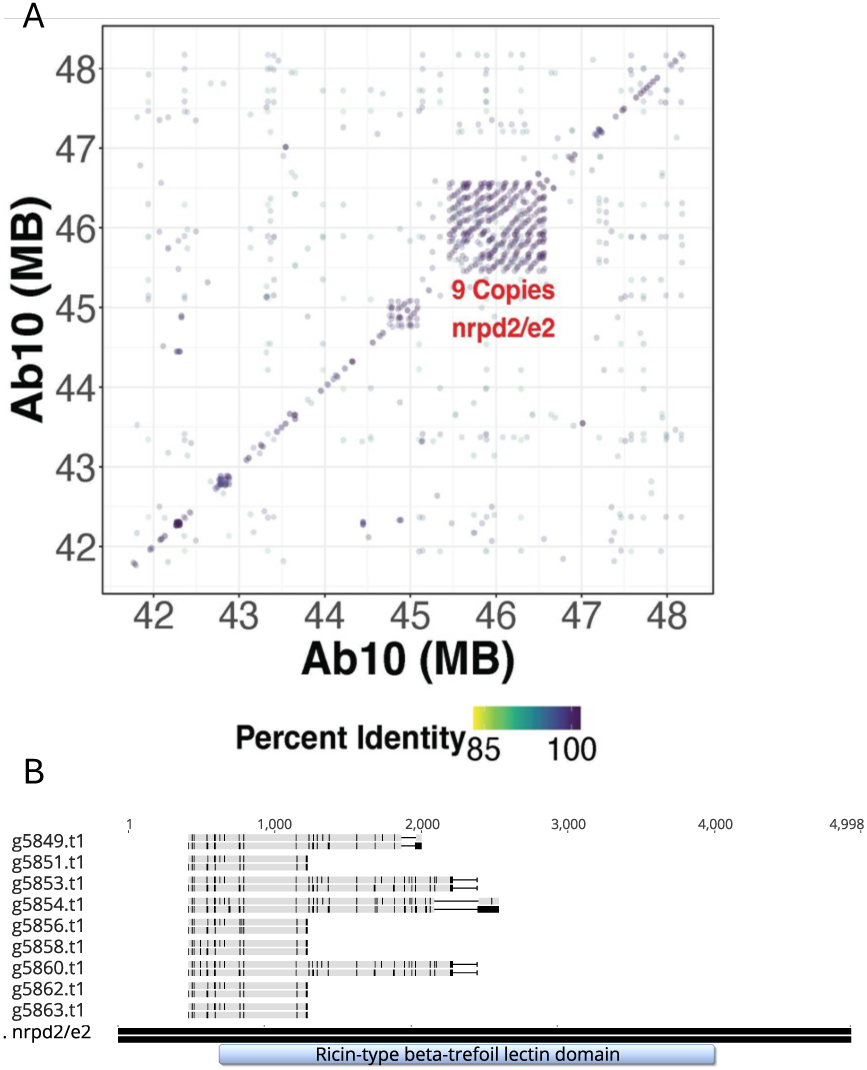
Subset of Sequence Comparison of Ab10 to Ab10. A. Each dot marks the start of a maximal unique match (MUM) of at least 300 bp long between various chromosome 10 haplotypes all of which begin at the *colored1* gene (Marçais *et al*. 2018). The color of each dot represents the percent identity of that match. All large knob arrays were removed for the sake of clarity. Ab10 refers to the assembly generated in this work. Array with 9 copies of the nrpd2/e2 homolog is marked. B. Alignment of the coding sequence of the 9 copies of the nrpd2/e2 homolog to the coding sequence of their closest homolog with the only protein functional domain marked. For all genes the top bar indicates the DNA coding sequence and the bottom line represents the protein translation. Grey indicates sequence that is identical to the nrpd2/e2 reference and black indicates sequence that is different from nrpd2/e2 the reference.

**Supplementary Figure 4:**
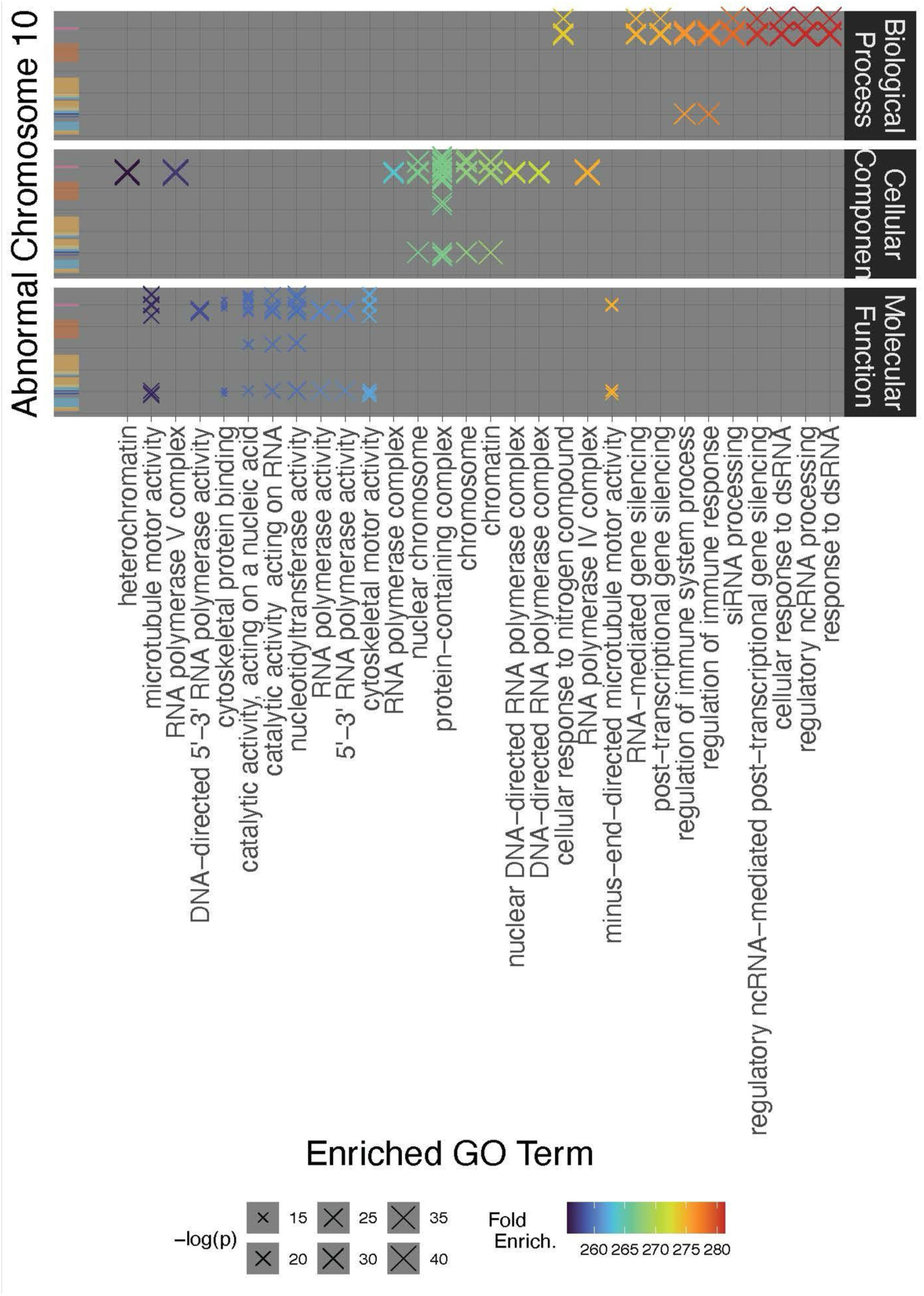
GO term enrichment on Ab10 non-shared Regions. Non-shared region refers to the regions with no consistent homology to normal chromosome 10. The y axis represents significantly enriched GO terms, the x axis indicates where genes associated with that GO term are located on the Ab10 haplotype. Color of each X represents fold enrichment, size represents statistical significance of enrichment. Relevant regions are marked by colored boxes: gold = shared, light blue = TR1 knob, dark blue = *trkin*, dark orange = knob180 knob, pink = *kindr*.

**Supplementary Figure 5:**
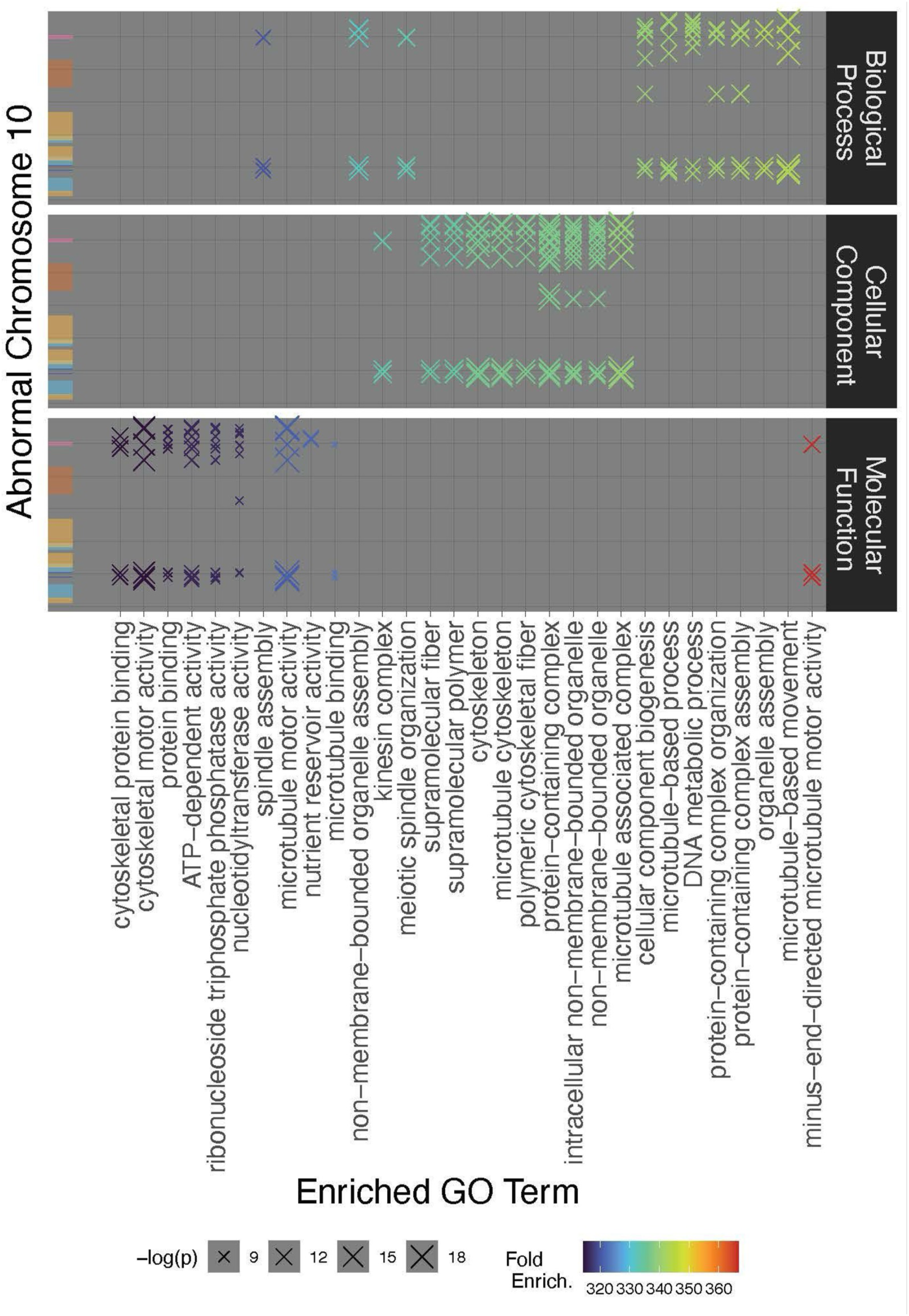
GO term enrichment on Ab10 non-shared Regions Without Duplicate Genes. Non-shared region refers to the regions with no consistent homology to normal chromosome 10. The y axis represents significantly enriched GO terms, the x axis indicates where genes associated with that GO term are located on the Ab10 haplotype. Color of each X represents fold enrichment, size represents statistical significance of enrichment. Relevant regions are marked by colored boxes: gold = shared, light blue = TR1 knob, dark blue = *trkin*, dark orange = knob180 knob, pink = *kindr*.

**Supplementary Figure 6:**
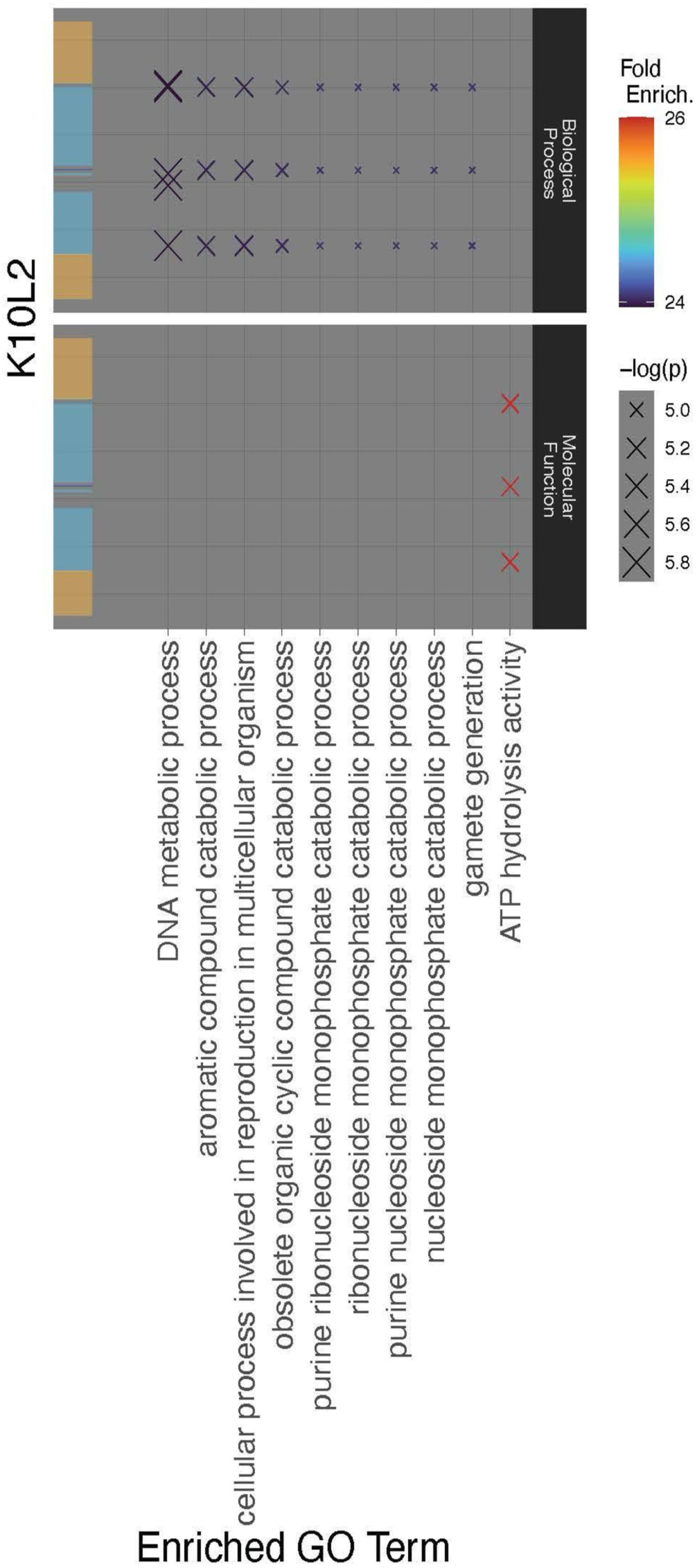
GO term enrichment on K10L2 non-shared Regions. Non-shared region refers to the regions with no consistent homology to normal chromosome 10. The y axis represents significantly enriched GO terms, the x axis indicates where genes associated with that GO term are located on the K10L2 haplotype. Color of each X represents fold enrichment, size represents statistical significance of enrichment. Relevant regions are marked by colored boxes: gold = shared, light blue = TR1 knob, dark blue = *trkin*.

**Supplementary Figure 7:**
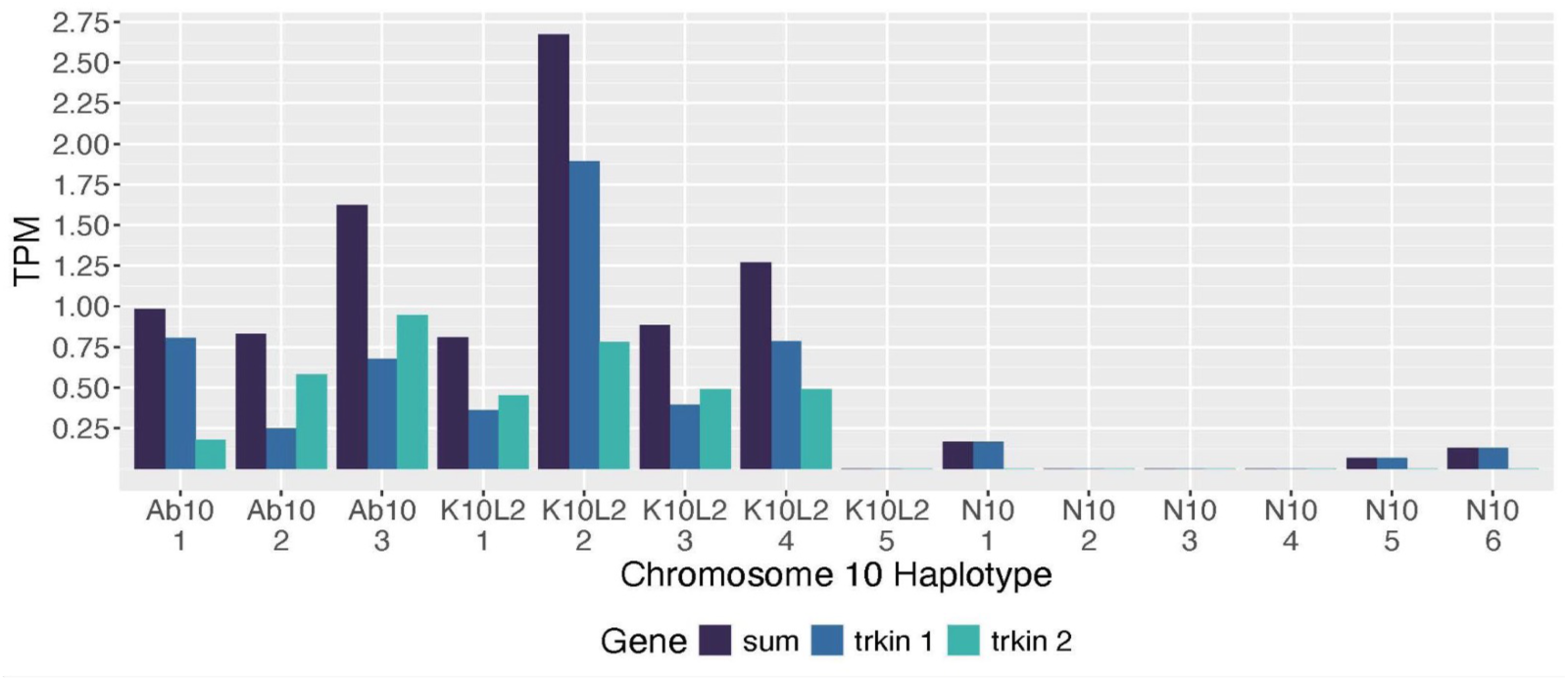
Expression of *Trkin* in Ab10, K10L2, and N10. Transcripts per million (TPM) for Ab10 *Trkin1* and *Trkin2* as well as their sum from mRNA sequencing data (Swentowsky *et al*. 2020).

**Supplementary Figure 8.**
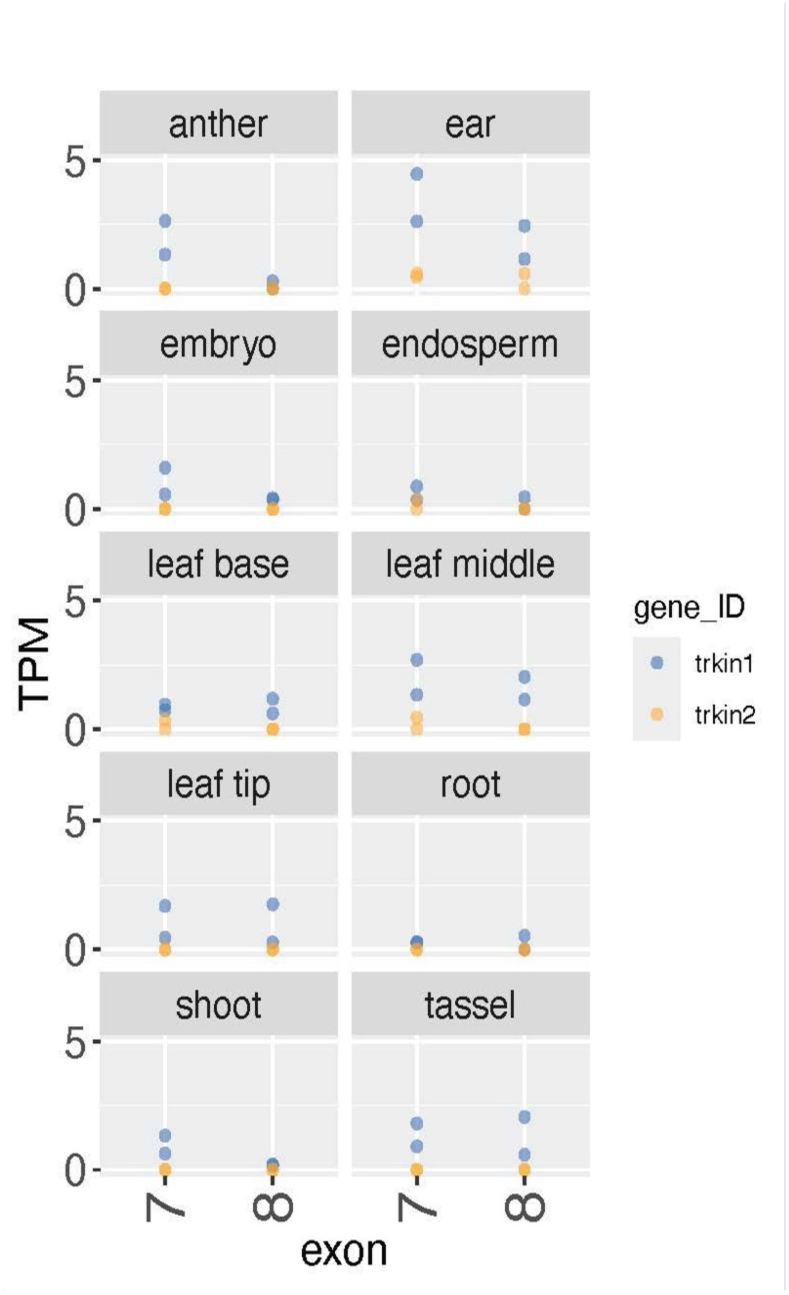
Expression of Ab10 *Trkin1* and *Trkin2*. TPM indicates transcripts per million. Each tissue was sequenced in two replicates indicated by two points per gene (color). Only exons 7 and 8 are differentiable between Ab10 Trkin1 and Trkin2, so only those were compared. Ab10 Trkin 1 mean expression was 1.078 TPM and Ab10 trkin 2 mean expression was 0.069 TPM. Welch t sample t test revealed the expression of exons 7 and 8 were different between Ab10 *Trkin1* and Ab10 *Trkin2* were significantly different (t = 6.5734, df = 41.476, p-value = 6.286e-08).

**Supplementary Figure 9:**
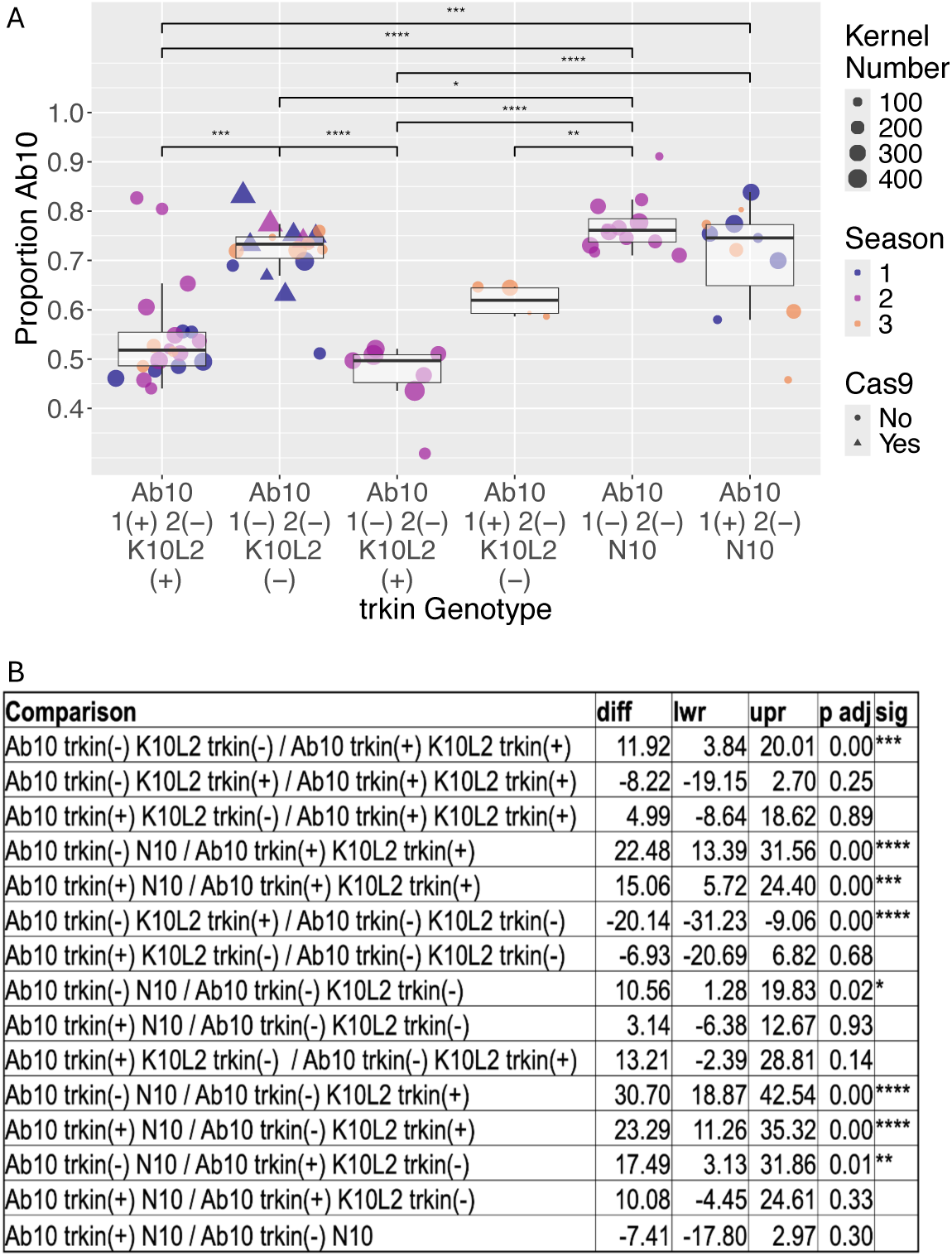
Effect of *Trkin* on K10L2 Ab10 Competition showing all comparisons and significance values. All plants were grown in the greenhouse in Athens, GA. Each dot represents an individual plant. Season refers to a group of plants grown at the same time. Seasons 1 and 2 were conducted in the same background while Season 3 was conducted in a different background. Season 1 and 2 of the Ab10 *trkin1(-) trkin2(-)* and *K10L2 trkin(-)* had *Cas9* segregating. The multi-way ANOVA model was Proportion Ab10 ∼ *Cas9* genotype + Round + *trkin* genotype. *Cas9* genotype = F(1,63)=9.656, p=0.00; Round = F(2,63)=0.520 p=0.59726; *trkin* genotype= F(5,63)=19.495, p= 1.11e-11. B. Results for Tukey’s HSD Test for multiple comparisons between all genotypes. diff=estimate of effect size, lwr = lower bound of 95% confidence interval, upr= upper bound of 95% confidence interval, p adj = p value adjusted for multiple comparisons, sig = symbol used. *=<0.05, **=<0.01, ***, <0.001, ***=0 *=<0.05, **=<0.01, ***, <0.001, ***=0. A. All significant relationships shown. B. Statistical output for all comparisons made.

**Supplementary Figure 10:**
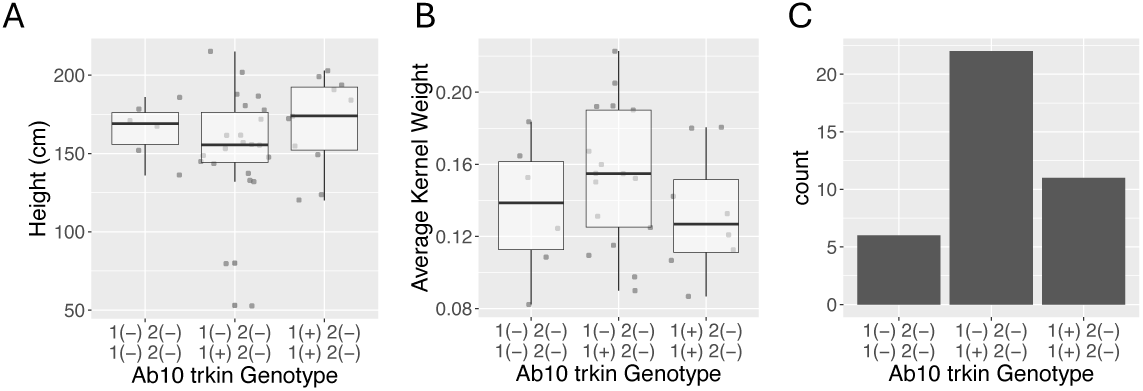
Plant fitness Effects of *trkin* in Ab10 Homozygotes. Plants were grown in the green house in Athens GA in a fully randomized order. A. Height: Fitted linear regression model was: Height ∼ Pot + Position in Greenhouse + *trkin* Genotype R^2^=0.1202, F(17,21)=1.305, p=0.2782. B. Average Kernel Weight: Fitted linear regression model was: Average Kernel Weight ∼ Pot + Position in Greenhouse + Silking time + Anthesis Time + *trkin* Genotype R^2^=0.3251 F(20,10)=1.723, p=0.1891. D,E. C. Transmission: Chi squared test was used to determine if the observed segregation of the *trkin* genotypes fit with Mendelian segregation X-squared(2)=1.2671, p=0.5307.

**Supplementary Figure 11:**
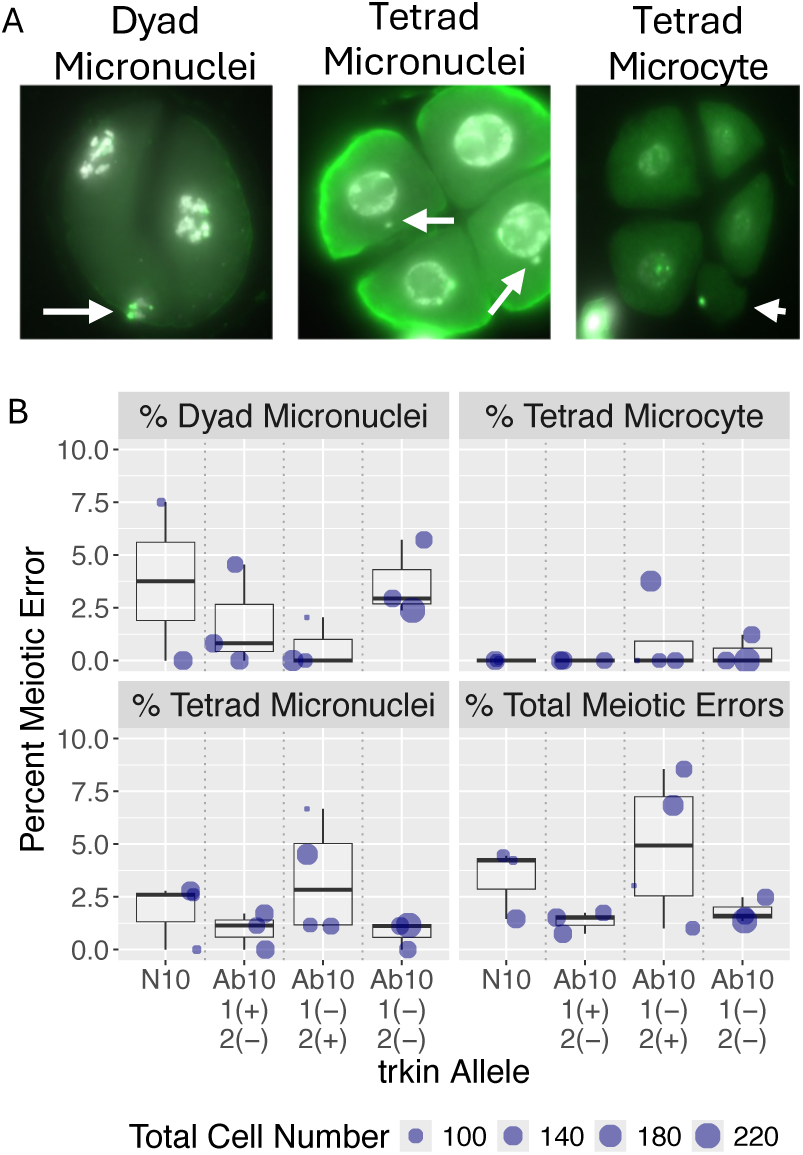
Meiotic Errors in Male Meiocytes of Various *Trkin* Genotypes. Meiotic errors were scored on Ab10 or N10 homozygous plants stained for FISH (Figure 7) with the indicated *trkin* genotypes. Dyad micronuclei refers to a lost chromosome at the conclusion of meiosis I. Tetrad micronuclei refers to a lost chromosome at the conclusion of meiosis II. Tetrad microcyte refers to an additional small cell containing DNA likely representing a lost chromosome at the conclusion of meiosis II. A. Contains examples of all scored meiotic errors. Cells with meiotic errors were normalized against the total number of same stage cells observed. Each dot represents an individual plant. One Way ANOVA determined no statistical difference in any class of meiotic error between *trkin* genotypes: % Dyad Micronuclei (F(3,9)=0.413, p= 0.748, % Tetrad Micronuclei (F(3,9)=1.552, p= 0.268, % Tetrad Microcyte (F(3,9)=0.549, p= 0.661, % Total Meiotic Errors (F(3,9)=1.89 p= 0.202.

**Supplementary Figure 12:**
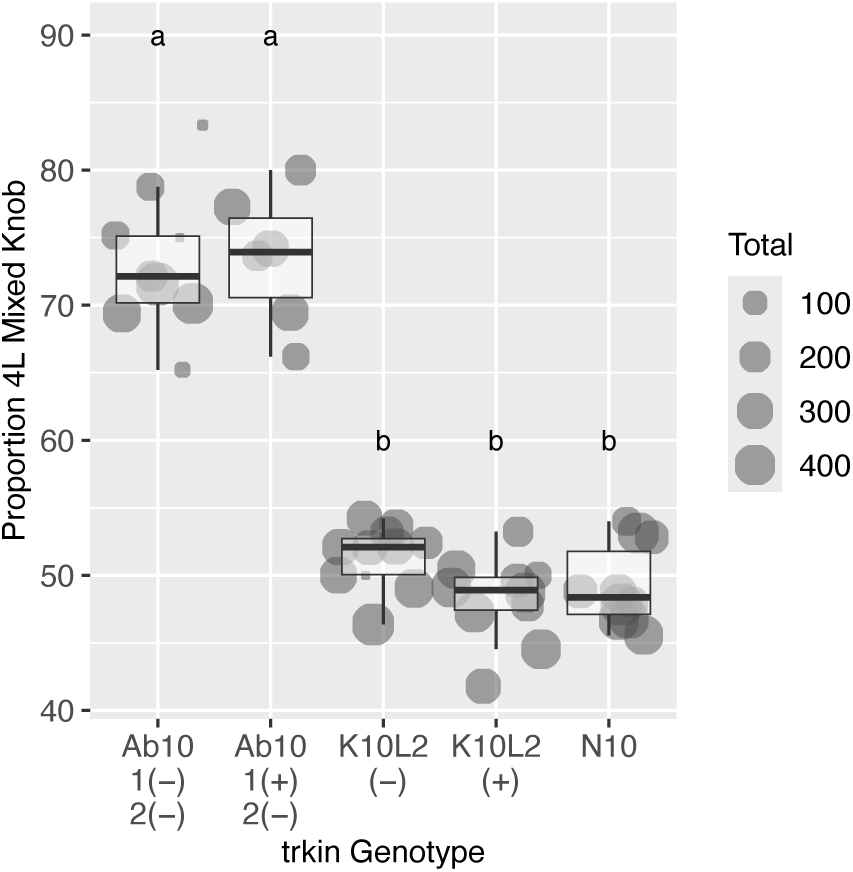
Effect of *Trkin* on Segregation of a Mixed Knob Not on Chromosome 10. All plants were grown in the greenhouse in Athens, GA. Each dot represents an individual plant. One way ANOVA model was Proportion 4L Mixed Knob ∼ *trkin* genotype F(4,41)= 101, p=<2e-16. Tukey’s HSD Test for multiple comparisons found that the mean value of all Ab10 bearing lines were significantly different from all K10L2 and N10 bearing lines (all p values = 0.00). Ab10 lines were not significantly different from each other. K10L2 and N10 lines were not significantly different from each other.

**Supplementary Table 1:**
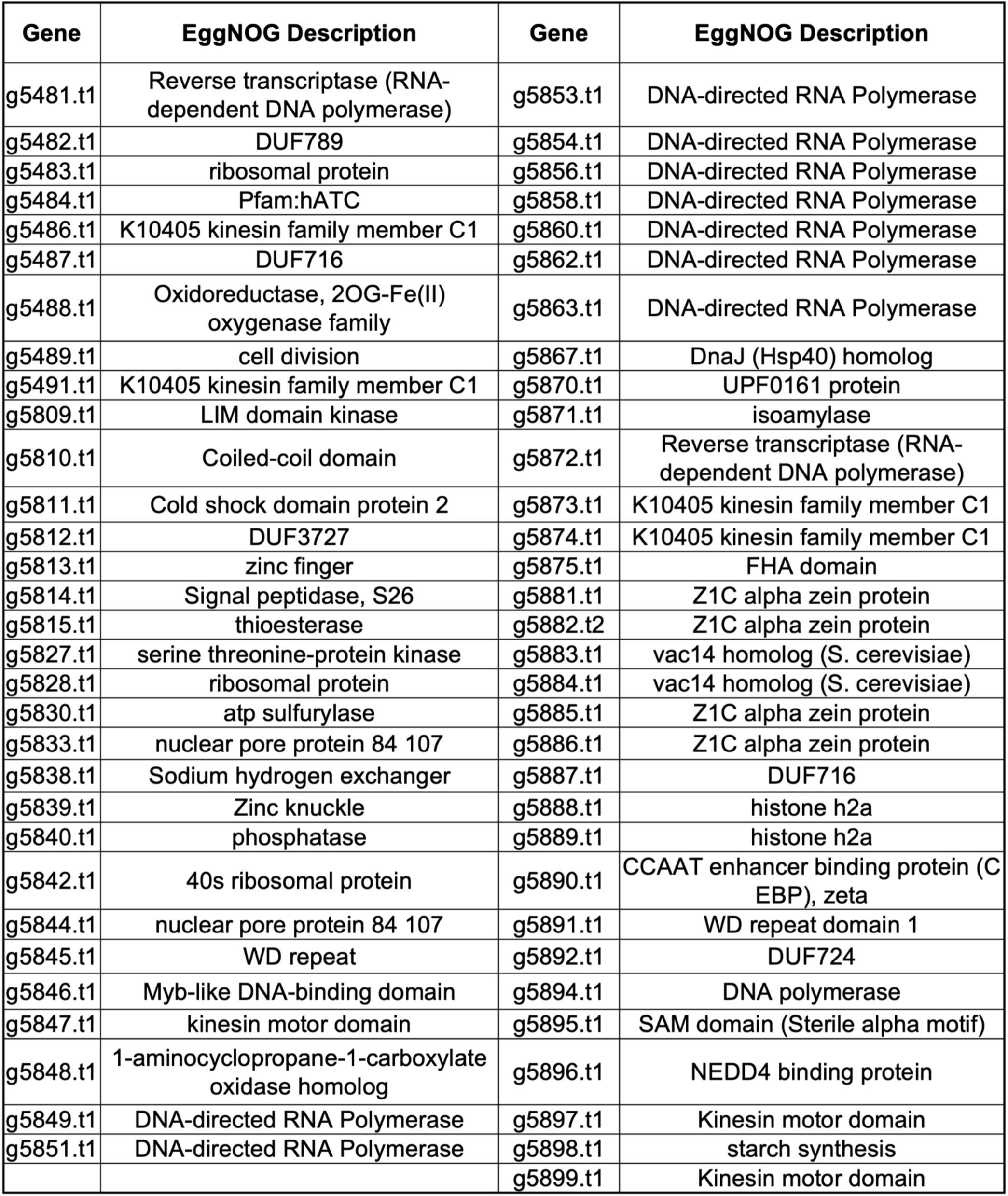
Ab10 non-shared genes functional annotation. Only genes with an EggNOG description are shown.

**Supplementary Table 2:** K10L2 non-shared genes functional annotation. Only genes with an EggNOG description are shown.

| **Gene** | **EggNOG Description** |
| --- | --- |
| g246.t1 | Multicopper oxidase |
| g247.t1 | complex ard1 subunit |
| g250.t1 | Reverse transcriptase (RNA-dependent DNA polymerase) |
| g254.t1 | Nicotiana lesion-inducing like |
| g255.t2 | FES |
| g256.t2 | Oxidoreductase, 2OG-Fe(II) oxygenase family |
| g257.t1 | cell division |
| g258.t1 | K10405 kinesin family member C1 |
| g262.t1 | Pentatricopeptide repeat-containing protein |
| g263.t1 | complex ard1 subunit |
| g264.t1 | Tyrosine 3-monooxygenase tryptophan 5-monooxygenase activation protein |
| g266.t1 | Coiled-coil domain-containing protein |
| g267.t1 | Protein of unknown function (DUF674) |
| g268.t1 | HLH |
| g269.t2 | Cold shock domain protein 2-like protein |
| g270.t1 | organic cation |
| g272.t1 | NmrA-like family |

**Supplementary Table 3:** Gene ortholog pairs on the non-shared regions of both Ab10 and K10L2.

| **Ab10 ID** | **K10L2 ID** | **EggNOG Description** |
| --- | --- | --- |
| g5475 | gA5475 | NA |
| g5479 | g251 | NA |
| g5486 | g258 | K10405 kinesin family member C1 |
| g5487 | gA5487 | Family of unknown function (DUF716) |
| g5488 | g256 | Oxidoreductase, 2OG-Fe(II) oxygenase family |
| g5489 | g257 | cell division |
| g5491 | g258 | K10405 kinesin family member C1 |
| g5847 | g258 | kinesin motor domain containing protein |
| g5867 | g269 | DnaJ (Hsp40) homolog, subfamily B, member 11 |
| g5873 | g258 | K10405 kinesin family member C1 |
| g5874 | g258 | K10405 kinesin family member C1 |
| gK255 | g255 | FES |

**Supplementary Table 4:**
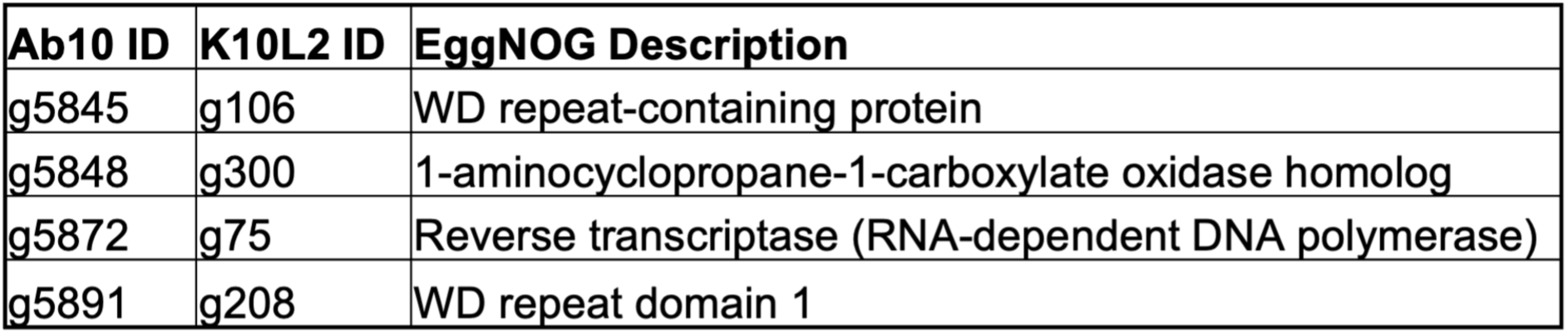
Gene ortholog pairs on the shared region of K10L2 and the non-shared region of Ab10.

| **Ab10 ID** | **K10L2 ID** | **EggNOG Description** |
| --- | --- | --- |
| g5845 | g106 | WD repeat-containing protein |
| g5848 | g300 | 1-aminocyclopropane-1-carboxylate oxidase homolog |
| g5872 | g75 | Reverse transcriptase (RNA-dependent DNA polymerase) |
| g5891 | g208 | WD repeat domain 1 |

**Supplementary Table 5:** Gene ortholog pairs on N10 and the non-shared region of Ab10.

| **Ab10 ID** | **B73 ID** | **EggNOG Description** |
| --- | --- | --- |
| g5820 | Zm00001eb430320 | NA |
| g5849 | Zm00001eb431870 | DNA-directed RNA Polymerase |
| g5851 | Zm00001eb431870 | DNA-directed RNA Polymerase |
| g5853 | Zm00001eb431870 | DNA-directed RNA Polymerase |
| g5854 | Zm00001eb431870 | DNA-directed RNA Polymerase |
| g5856 | Zm00001eb431870 | DNA-directed RNA Polymerase |
| g5858 | Zm00001eb431870 | DNA-directed RNA Polymerase |
| g5860 | Zm00001eb431870 | DNA-directed RNA Polymerase |
| g5862 | Zm00001eb431870 | DNA-directed RNA Polymerase |
| g5863 | Zm00001eb431870 | DNA-directed RNA Polymerase |

**Supplementary Table 6:**
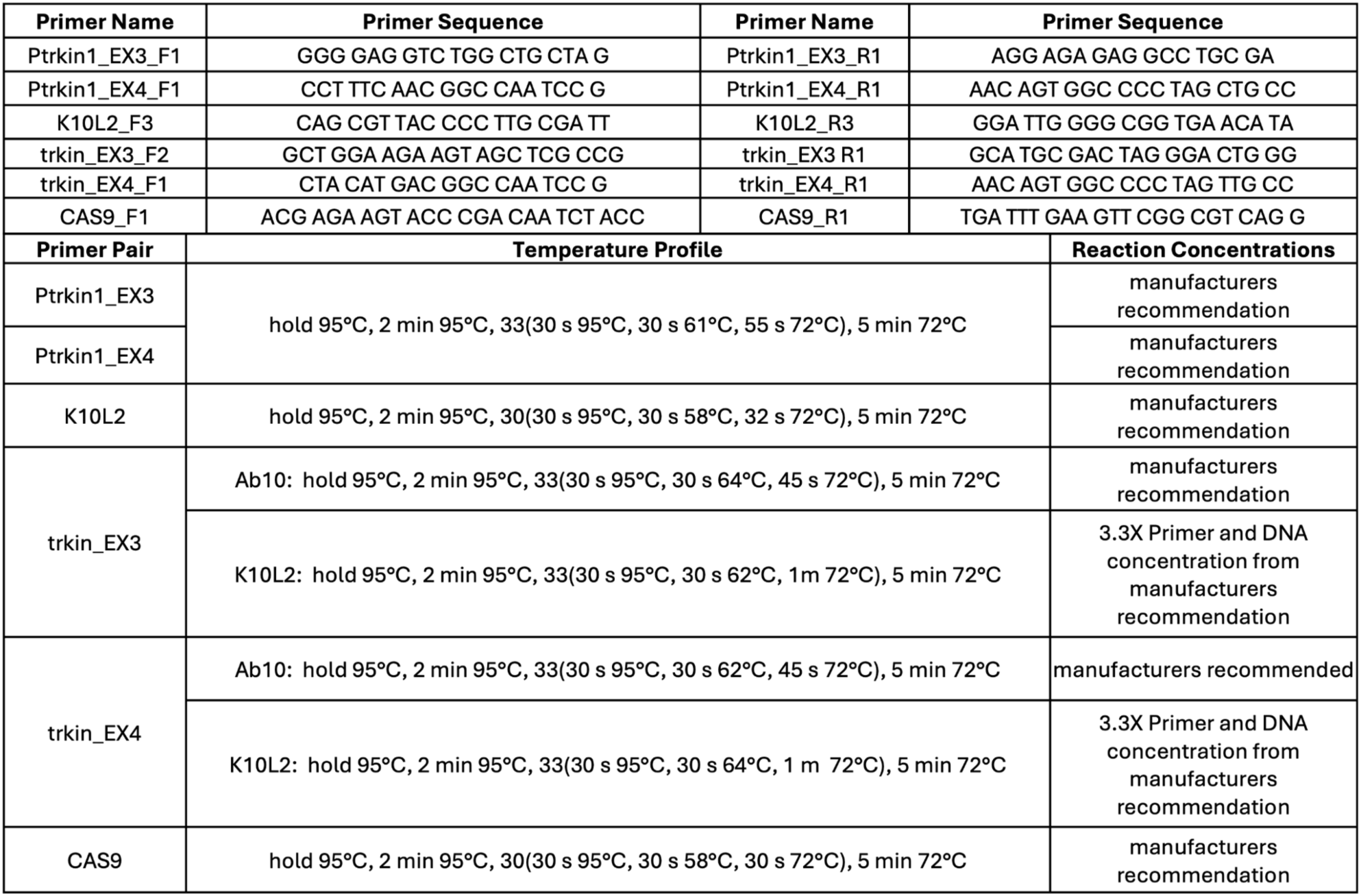
Primers and Reaction Parameters Used For Genotyping.

## DATA AVAILABILITY

All code, the Ab10 and K10L2 haplotype assemblies and genome annotations, and supplemental file 1 are available at https://github.com/dawelab/TRKIN_Published.git. Raw PacBio HiFi data for Ab10 and CI66 are being submitted to the NCBI SRA.

## ACKNOWLEDGMENTS

We thank Dr. Jianing Liu for her assistance in the B73-Ab10 v2 assembly. We thank Tanvi Kamat and Anne Blevins for sorting and counting kernels. We thank Dr. Zenglu Li and lab for generously allowing us to use their seed counter. We thank the Georgia Advanced Computing Resource Center for their technical expertise.

